# Exercise Engages Coordinated Neuron–Glia Signaling to Shape Spinal Cord Plasticity

**DOI:** 10.64898/2025.12.11.693787

**Authors:** Shivani Mansingh, Sedat Dilbaz, Danilo Ritz, Gesa Santos, Stefan A. Steurer, Christoph Handschin

**Affiliations:** Biozentrum, University of Basel, Basel, Switzerland

**Keywords:** spinal cord, exercise, glial plasticity, snRNA sequencing, neuron-glia communication

## Abstract

Physical activity induces systemic benefits for brain and muscle function, but how the healthy spinal cord adapts to exercise remains largely unknown. Here, we combine bulk proteomics, single-nucleus RNA sequencing, and cellular communication inference to map exercise-induced molecular adaptations in the mouse lumbar spinal cord. Endurance training elicited robust baseline remodeling, dominated by glial transcriptional changes. Acute exhaustive exercise triggered biphasic responses: widespread metabolic and synaptic gene upregulation at 6 h followed by balanced suppression at 24 h, with trained animals exhibiting enhanced amplitude and faster resolution. Cell–cell communication analysis revealed that exercise reshaped signaling networks in both magnitude and composition. While glia emerged as primary transcriptional responders, cholinergic neurons—despite minimal transcriptional changes—were central signaling hubs, engaging various pathways with astrocytes, oligodendrocyte precursor cells, and oligodendrocytes in a training-dependent and temporally restricted manner. Glial-derived communication further diversified these responses, with astrocytes, oligodendrocytes, and microglia shifting toward pathways supporting synaptic remodeling, axon guidance, and growth factor signaling while dampening inflammatory cues. Together, these findings identify neuron–glia communication as potential driver of spinal cord adaptation to exercise, suggesting pathways through which glial plasticity may serve as a key mediator linking motor activity to spinal cord resilience.

**Significance Statement:** Exercise confers broad benefits for neural and motor system health, yet how the healthy spinal cord adapts to physical activity is poorly understood. Using multi-omic profiling, we show that endurance training and acute exercise elicit coordinated transcriptional and signaling responses in spinal cord glia, with cholinergic neuron–glia communication emerging as a key upstream driver. These findings identify glial plasticity and neuron–glia crosstalk as a potential mechanism of spinal cord adaptation, providing a foundation for leveraging exercise-based strategies in neurological health and disease.

## INTRODUCTION

Physical exercise is a potent modulator of central nervous system plasticity, promoting structural and functional adaptations across diverse neuronal networks. While extensive research has characterized exercise-induced changes in the brain, particularly within hippocampal circuits (1–5), the spinal cord represents a remarkably understudied yet critical component of the exercise-neuroplasticity axis. The spinal cord integrates descending motor commands with ascending sensory information, positioning it to coordinate exercise-related neural adaptations (6).

The therapeutic potential of exercise-induced spinal plasticity has been predominantly studied in injury contexts. After spinal cord injury, exercise promotes neurotrophin expression, enhances synaptic transmission, and facilitates oligodendrocyte proliferation (7–13). Voluntary wheel running is particularly effective compared to forced paradigms, promoting enhanced neurotransmitter modulation and axonal innervation (12, 14–17). These studies implicate pathways such as brain-derived neurotrophic factor (BDNF) signaling, synaptic vesicle cycling, and glial-mediated responses in these processes (7–9, 11, 15, 17, 18). However, a critical knowledge gap exists regarding exercise effects in the healthy spinal cord. Understanding these foundational adaptations is essential for various reasons, e.g. to understand the consequences of increased and repeated (hyper-) activation of motor systems in exercise (often over hours and to failure), thereby revealing neuroprotective processes that may be leveraged in age- or disease-related degeneration.

A multicellular involvement in the response of the spinal cord to repeated bouts of exercise could be expected from prior data in the spinal cord as well as insights gained in the brain. Glial cells demonstrate exquisite sensitivity to activity-dependent stimuli (4, 10, 12, 16, 19–22). Microglia orchestrate neuroplasticity responses and coordinate cellular interactions during repair, with exercise promoting anti-inflammatory responses while suppressing pro-inflammatory activation, and playing additional roles, e.g. in mediating neuropathic pain (23–27). Astrocytes in the brain respond to physical activity through metabolic and structural adaptations that support neuronal function (28–33), but their specific responses in the healthy spinal cord remain largely unexplored (34). Oligodendrocytes are also highly exercise-sensitive, with voluntary running enhancing precursor proliferation through Wnt signaling and increasing nestin-positive progenitors in spinal white and gray matter (21). Motor skill learning specifically requires rapid oligodendrocyte production, and exercise promotes peroxisome proliferator-activated receptor γ coactivator 1α (PGC1α)-mediated myelin synthesis, suggesting these cells facilitate adaptive myelination in response to increased neural activity (12, 13).

Recent advances in single-nucleus RNA sequencing (snRNA-seq) have revolutionized our understanding of cellular heterogeneity within spinal cord circuits, revealing extensive glial diversity in the human and mouse spinal cords (35–44). These approaches overcome technical barriers of adult tissue dissociation and enable unbiased resolution of exercise-responsive populations. When integrated with proteomics, they provide comprehensive insight into both cell-type-specific transcriptional dynamics and tissue-wide protein-level adaptations. Two recent studies applied snRNA-seq to examine how aerobic exercise influences the aging spinal cord, showing that training can partially restore youthful cellular and regulatory features (45, 46). While these datasets advance our broader understanding of exercise–aging interactions, several features of exercise-induced adaptation in the healthy adult spinal cord remain insufficiently defined. In particular, neither study captured acute temporal dynamics following an exercise bout, assessed inter-individual variability due to pooled sequencing designs, or examined lumbar segments that directly innervate hindlimb musculature. Additionally, both relied on fresh-tissue dissociation, which is susceptible to stress-induced transcriptional artifacts that newer fixed-tissue approaches help to mitigate.In this study, we use multi-omic profiling to characterize how endurance training and acute exercise remodel the healthy mouse spinal cord. We combine voluntary wheel running with bulk proteomics and fixed-tissue snRNA-seq (10x Genomics Flex) to profile lumbar spinal cord segments at baseline and at 6 h and 24 h after an acute bout of exercise in both sedentary and trained mice. This design allows us to distinguish transient from long-lasting adaptations, quantify cell-type–specific transcriptional programs and signaling pathways with independent biological replicates, and focus particularly on glial plasticity and neuron–glia crosstalk. Our work therefore provides a temporally and anatomically resolved, multi-omic framework for exercise-induced adaptations in the young, healthy spinal cord.

## RESULTS

### Endurance Training Remodels Spinal Cord Composition and Glial Gene Expression

To investigate how prolonged physical activity impacts the spinal cord, we subjected adult male C57BL/6 mice to six weeks of voluntary wheel running (TRAIN), with sedentary littermates serving as controls (SED) (Figure 1A). Whole-body composition and exercise capacity were assessed before and after the intervention. Mice with access to running wheels voluntarily ran between 4 and 8 km/day, averaging approximately 6 km/day over the training period (Figure 1B). While total body mass increased in both groups (Figure S1A), trained mice preserved lean mass, whereas sedentary mice showed significant decreases in relative lean mass combined with an increase in fat mass (Figure S1B–C). Treadmill-based endurance testing demonstrated an approximately threefold increase in maximal exercise capacity in trained mice (Figure 1C), validating the effectiveness of our aerobic training protocol.

**Figure 1.**
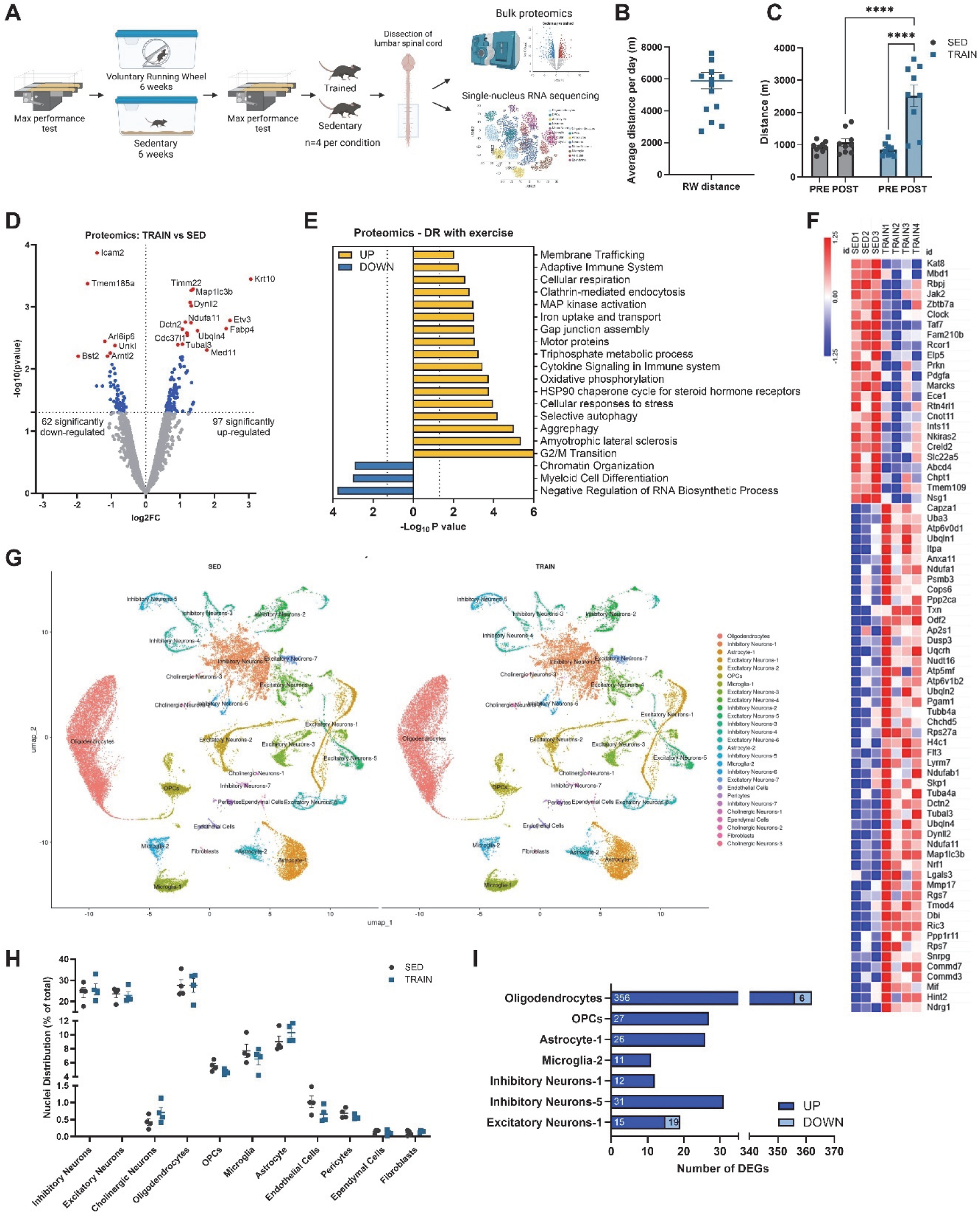
Endurance Exercise Induces Molecular and Cellular Changes in Adult Mouse Spinal Cord. (A) Experimental design: Adult male C57BL/6 mice underwent six weeks of voluntary wheel running (TRAIN) or remained sedentary (SED), followed by performance testing and spinal cord dissection (n=15 per condition). Then a subset of the cohort (n=4 per condition) was used for bulk proteomics and single-nucleus RNA sequencing (snRNA-seq). (B) Average daily running distances of TRAIN mice over the intervention period. Each data point reflects an individual mouse; error bars denote mean ± SEM (n = 16 mice). (C) Treadmill-based endurance performance measured pre- and post-intervention in SED and TRAIN groups. Bars depict mean ± SEM; individual data points shown. ****p < 0.0001, two-way ANOVA with multiple comparisons (Fisher’s LSD test). (D) Volcano plot: Differentially expressed proteins in lumbar spinal cord (TRAIN vs. SED); proteins with p < 0.05 and |log₂FC| >= 0.25 are highlighted in blue and top differentially regulated (DR) proteins in red. (E) Top Gene Ontology (GO) terms for DR proteins: upregulated (yellow), downregulated (blue). (F) Heatmap of all significant (p < 0.05 and |log₂FC| >= 0.5) differentially expressed proteins (scaled by z-score). (G) UMAP visualization of major spinal cord cell populations from snRNA-seq, colored by cell identity and split by condition (SED, TRAIN). (H) Quantification of cell-type abundance (percent of total nuclei) by group. Bars depict mean ± SEM; n=4 per group, individual data points shown. No significant changes observed; two-way ANOVA with Šídák’s multiple comparisons test. (I) Bar plot showing number of differentially expressed genes (DEGs, up/down) in each main cell-type after training. Genes were filtered for |log₂FC| > 0.5 and q < 0.01. Abbreviations: SED, sedentary; TRAIN, voluntary exercise group; UMAP, Uniform Manifold Approximation and Projection; FC, fold-change; SEM, standard error of the mean.

To determine whether the endurance training paradigm induces molecular remodeling in the spinal cord at rest, i.e. untainted by the last acute exercise bout, we performed quantitative bulk proteomics on lumbar spinal cord lysates. 159 proteins were differentially expressed between trained and sedentary mice (p < 0.05, |log₂FC| > 0.5); including 97 upregulated and 62 downregulated proteins (Figure 1D). Gene Ontology (GO) enrichment analysis revealed upregulated proteins to be linked to mitochondrial metabolism, vesicle-mediated transport, autophagy, and stress response pathways, whereas downregulated proteins were enriched for chromatin organization and immune-related functions (Figure 1E). Top upregulated proteins—including TIMM22, NDUFA11, UBQLN4, MAP1LC3B, DYNLL2, TUBB4A, NRF1, ETV3, and FABP4 —reflect regulation of neurodevelopmental signaling, chromatin remodeling, cytoskeletal and metabolic adaptation, and CNS homeostasis (Figure 1F).

To resolve the cellular origin of these proteomic changes, we performed high-throughput single-nucleus RNA sequencing (snRNA-seq) using the 10x Genomics Fixed RNA Profiling (Flex) protocol (Figure 1A, S1D-E). This method enabled nuclei isolation from frozen adult central nervous system (CNS) tissue, providing improved recovery of intact transcriptional profiles, with enhanced sensitivity for low-abundance transcripts and greater uniformity across nuclei. Using canonical gene markers, we annotated 11 major cell-types, including neurons, astrocytes, oligodendrocyte lineage cells, microglia, vascular cells, and ependymal cells, with finer substructure within multiple glial populations (Figure 1G). Estimates of cell-type abundance remained broadly stable between groups (Figure 1H), indicating that exercise training does not alter gross cellular composition in a persistent manner. Nevertheless, transcriptional reprogramming was evident, with glial cells—particularly oligodendrocytes— showing the most prominent differential gene expression in response to training (Figure 1I). These data suggest glial populations play a central role in mediating spinal adaptations to chronic physical activity.

### Glia are the Primary Transcriptomic Responders to Training

To identify the key cellular contributors to spinal cord remodeling post-endurance training, we focused on glial populations showing the most pronounced transcriptional changes: oligodendrocytes, oligodendrocyte precursor cells (OPCs), and the Astrocyte-1 subcluster (Figure 2A–C). Among these, oligodendrocytes had the greatest number of differentially expressed genes (DEGs). Gene set enrichment analysis (GSEA) indicated that shared biological processes—such as synaptic modulation, axon development, and intercellular signaling— were activated across these glial populations (Figure 2D). However, the specific genes driving these enrichments were largely cell-type specific, with only minimal DEG overlap (Figure S2A), except for the shared genes between oligodendrocytes and OPCs, consistent with their developmental relationship.

**Figure 2.**
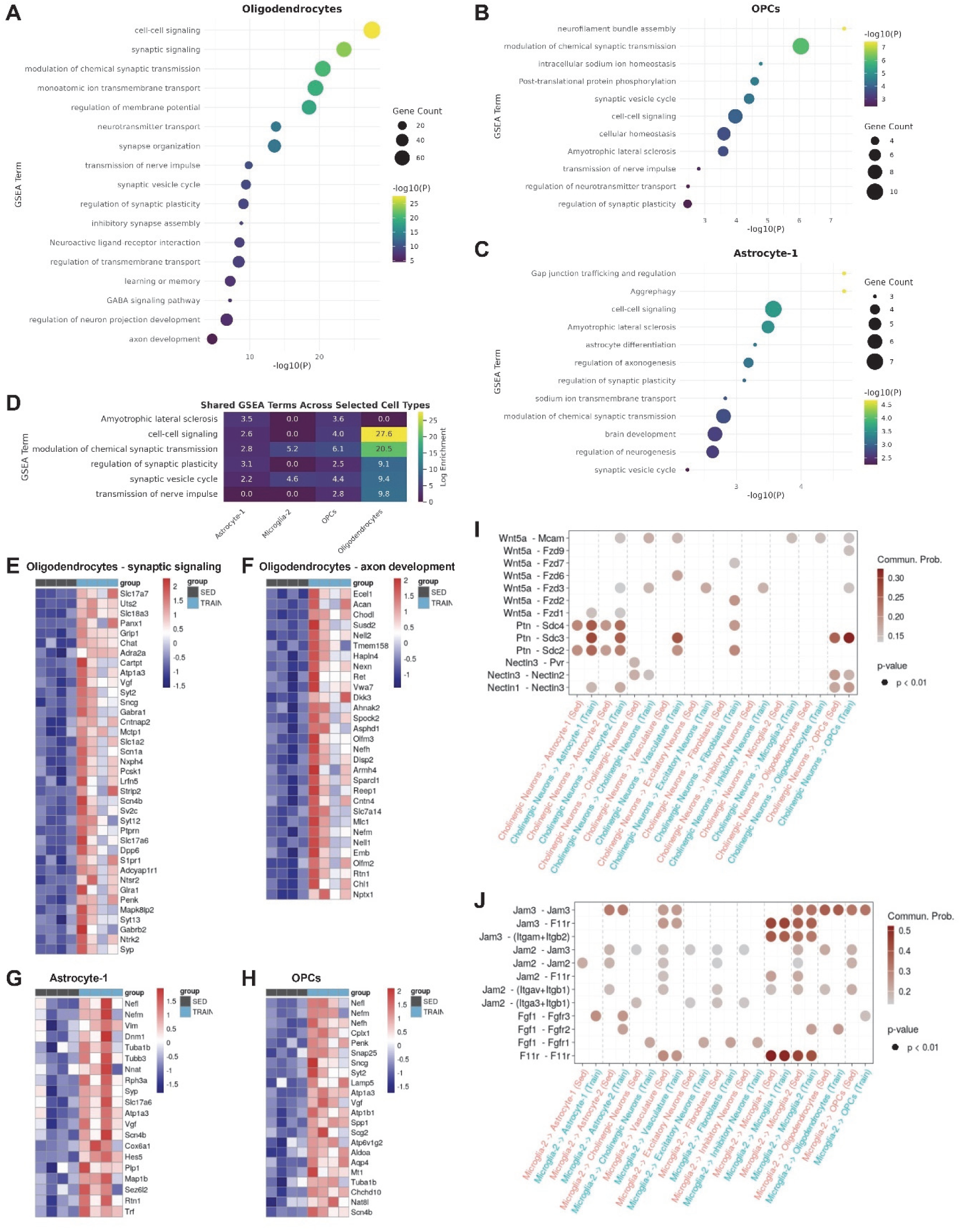
Glial Populations Mediate Cell-Type–Specific Transcriptional Remodeling and Intercellular Network Reorganization After Endurance Exercise. (A–C) Dot-plot of top gene set enrichment analysis (GSEA) terms associated with the differentially expressed genes up-regulated in trained spinal cords for oligodendrocytes (A), oligodendrocyte precursor cells (OPCs; B), and Astrocyte-1 (C) clusters. (D) Heatmap summarizes the overlap of key GSEA terms across selected glial cell-types after exercise training, highlighting shared pathways of neural adaptation. (E–F) Heatmaps illustrate expression of representative genes upregulated in oligodendrocytes, grouped by synaptic signaling (E) and axon development (F). Expression levels are indicated by color (red = upregulated, blue = downregulated) and grouped by sedentary (SED) or trained (TRAIN) state. All genes displayed were filtered for |log₂FC| > 0.5 and q < 0.01. (G–H) Heatmaps show top upregulated genes in Astrocyte-1 (G) and OPCs (H) after training, with relative expression indicated by color scale across groups. (I–J) Dot plots depict communication probabilities for selected ligand–receptor interactions: WNT, Pleiotrophin (PTN), and Nectin signaling from cholinergic neurons to various spinal cord cell-types (I), and JAM and FGF signaling from microglia-2 to all spinal cord populations (J), under sedentary (SED) and trained (TRAIN) conditions. Dot color represents communication probability all dots reflect significant interactions (p < 0.01).

In response to training, oligodendrocytes upregulated genes associated with vesicle trafficking and synaptic function (*Slc17a7, Syt2, Snap25, Vgf*), ion transport and membrane potential (*Atp1a3, Slc4a4, Panx1*), and axon-glia interactions (*Ret, Dkk3, Sparcl1, Hapln4*) (Figure 2E–F, S2B–C). Increased expression of metabolic and structural genes (*Acsl6, Mlc1, Reln, Slc24a4*) suggests enhanced activity-dependent support of axons. Trained OPCs were enriched for genes related to neurofilament organization, intracellular signaling, and synaptic modulation (Figure 2B), with upregulation of synaptic, metabolic, and structural genes (Figure 2G), consistent with emerging roles for OPCs in myelin plasticity and neuronal responsiveness (47, 48).

Astrocyte-1 displayed enriched programs related to gap junction regulation, cell–cell communication, and axonogenesis (Figure 2C), with significant upregulation of cytoskeletal and signaling regulators (*Vim, Rtn1, Map1b, Hes5, Trf, Sparcl1*) in trained compared to untrained spinal cord (Figure 2H). Although several DEGs (*Gja1, Syp, Vgf, Nefh*) were expressed in multiple glial subtypes, their expression trajectories differed, suggesting functionally distinct roles. These findings demonstrate that while glial subtypes converge on certain biological processes, their transcriptomic responses are highly cell-type specific and aligned with their specialized roles in the spinal cord.

Expanding the analysis to other cell-types, we found that microglia-2 displayed training-induced expression changes in genes associated with synaptic regulation and secretory function (*Dnm1, Snap25, Stxbp1, Sparcl1, Nrxn1*), as well as enriched GSEA terms for gliogenesis and synaptic modulation (Figure S2D, H). These data support a shift away from classical immune activation toward a more synaptically engaged, neuroprotective microglial phenotype.

By contrast, neuronal responses were more heterogeneous and generally of smaller magnitude. Excitatory neurons (ExN-1) showed modest upregulation of ion transport and homeostasis genes (*Slc1a2, Atp1a2, Slc4a4, Gja1*) and downregulation of immune-responsive genes (*Nr4a1, Ly6a*), suggesting reduced inflammatory signaling after training (Figure S2E, I). Inhibitory neuron subclusters responded variably: Inh-1 displayed upregulation of genes related to neuronal growth and cytoskeletal organization, while Inh-5 was enriched for genes in MAPK signaling, membrane transport, and neurotransmitter uptake (Figure S2F–G). Genes upregulated included *Agt, Apoe, Gja1, Uts2, Mt1, Mt3*, and various solute carrier transporters (Figure S2J–K), reflecting increased inhibitory tone and metabolic support.

### Endurance Training Reshapes Intercellular Signaling in the Spinal Cord

To determine if transcriptional changes were accompanied by altered intercellular communication, we used CellChat (49) to infer ligand–receptor (L–R) interactions among spinal cord cell-types. Endurance training remodeled multiple signaling networks, suppressing stress- and inflammation-associated pathways such as neurotensin (NTS), leukemia inhibitory factor receptor (LIFR), and carcinoembryonic antigen-related cell adhesion molecule (CEACAM) signaling, while enhancing activity in pathways including Notch, non-canonical WNT (ncWNT), and Galectin (Figure S2L). Notably, the predicted communication patterns did not simply mirror the magnitude of differential expression within each cell-type: neurons (inhibitory, cholinergic, excitatory) and supportive cells (fibroblasts, astrocytes, vasculature) emerged as major senders, while astrocytes, neurons, and OPCs were the predominant receivers (Figure 1I, S2L–M). Thus, even populations with relatively few DEGs contributed disproportionately to signaling networks, highlighting that exercise-induced plasticity involves a reorganization of intercellular communication beyond what is captured by cell-intrinsic transcriptional changes alone.

Further analysis of cell-type-specific L–R usage across key signaling pathways (Figure 2I–J, S3) confirmed that several communication axes were enhanced by training. WNT, PTN, and NECTIN signaling from cholinergic neurons to glial and vascular targets were mainly observed only in trained animals (Figure S2I), suggesting recruitment of additional structural and trophic signaling cues. Likewise, NRXN-DAG1 interactions between cholinergic neurons and astrocytes were strengthened by exercise training, potentially enhancing synaptic formation, stability, and communication (Figure S3A). NRXN signaling from astrocytes to neurons, OPCs, and other astrocytes also increased (Figure S3B), indicating broader post-training engagement of astrocyte-mediated synaptic and glial networks.

Laminin signaling derived from astrocytes and oligodendrocytes increased toward neuronal, vascular, and fibroblast targets with exercise (Figure S3C–D), consistent with remodeling of extracellular matrix and tissue architecture. Microglia-2 in trained mice exhibited reduced JAM-mediated signaling and amplified Semaphorin (SEMA4) and FGF signaling to several targets (Figure 2J, S3E), marking a possible transition from immune surveillance toward growth- and guidance-related signaling. Together, these findings show that physical activity drives marked, cell-type-specific remodeling of spinal cord intercellular communication networks.

### Acute Exercise Induces Distinct Temporal and Cell-Specific Molecular Responses Shaped by Training

Given the extensive transcriptional and proteomic remodeling evoked by endurance training at rest, we next sought to determine how the spinal cord temporally responds to a single acute bout of exhaustive exercise—and whether this response is modulated by prior training. Sedentary (SED) and trained (TRAIN) mice underwent a single session of treadmill-based running to exhaustion before lumbar spinal cords were collected at 6-hours (6h) and 24-hours (24h) post-exercise (Figure 3A).

**Figure 3.**
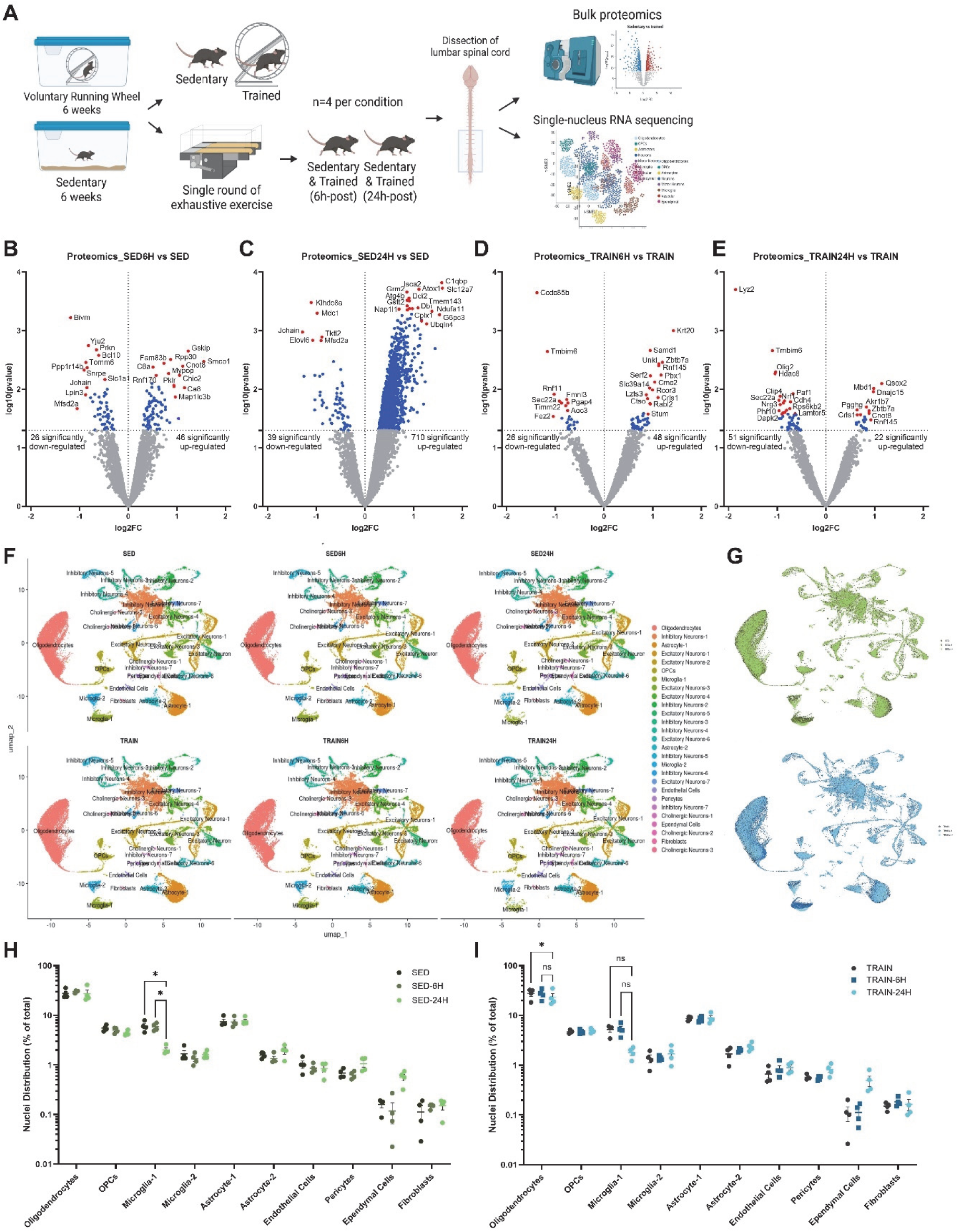
Acute Exhaustive Exercise Induces Distinct Temporal and Training-Dependent Proteomic and Cellular Remodeling in Spinal Cord. (A) Experimental design: Sedentary (SED) and endurance-trained (TRAIN) mice subjected to a single session of treadmill-based exhaustion exercise, with lumbar spinal cords collected at 6- and 24-hours post-exercise; n = 4 mice per group/timepoint. (B–E) Volcano plots from bulk proteomics showing differentially expressed proteins (DEPs) following exercise. (B) SED 6h vs. SED baseline, (C) SED 24h vs. SED baseline, (D) TRAIN 6h vs. TRAIN baseline, (E) TRAIN 24h vs. TRAIN baseline. Proteins with p < 0.05 and |log₂FC| >= 0.25 are highlighted in blue and top differentially regulated proteins in red. Numbers reflect total significantly up- or downregulated proteins per comparison (p < 0.05, |log₂FC| > 0. 5). (F) UMAP plots of integrated single-nucleus RNA sequencing data illustrate clustering and abundance of major spinal cord cell-types split across all six experimental conditions. (G) Spatial overlays of nuclei from each condition on the integrated UMAP, with green representing SED and blue representing TRAIN groups, to visualize cluster distribution and identify cell populations that diverge in UMAP space across conditions. (H–I) Quantification of non-neuronal cell-type proportions across conditions: (H) SED baseline, 6h, 24h; (I) TRAIN baseline, 6h, 24h. Proportions plotted as percent of total nuclei (mean ± SEM, n = 4/group). * p < 0.05, ns = not significant, 0.05<p<0.1 (two-way ANOVA with Tukey’s multiple comparisons test).

Bulk proteomics revealed a dynamic molecular response in sedentary mice, with modest changes at 6h and robust upregulation of 710 proteins by 24h (Figure 3B–D). In contrast, trained mice exhibited blunted proteomic responses at both time points, with fewer differentially expressed proteins and smaller fold changes (Figure 3D–E), implying that training buffers the acute molecular response and fosters more robust homeostatic control. Gene ontology enrichment analysis supported this interpretation. In sedentary mice, acute exercise strongly activated immune-related, metabolic, and mitochondrial pathways—particularly at 24h (Figure S4A–B). In trained animals, these same pathways were suppressed (Figure S4C–D), consistent with a shift from reactive to regulated responses.

To resolve the cellular contributors to these bulk changes, we performed single-nucleus RNA sequencing (snRNA-seq) on all six experimental conditions. Integration with prior datasets identified 11 major cell-types with stable clustering across samples (Figure 3F). Overall cell-type proportions remained largely unchanged, but there were subtle, dynamic shifts. Notably, Microglia-1 abundance decreased 24h post-exercise in sedentary animals, with a milder trend in trained mice (Figure 3H–I). Spatial overlays of nuclei suggest this reduction stems from a shift toward Microglia-2, potentially representing altered microglial activation (Figure 3G). Trained mice displayed a modest increase in oligodendrocytes at 24h and both groups showed a minor increase in inhibitory neuron subcluster 1 (Inh-1) after exercise (Figure 3G–I, S4F–G). These changes were absent at baseline, indicating they likely represent transient, exercise-induced transcriptomic shifts rather than chronic adaptive effects.

These data show that acute exhaustive exercise triggers rapid, extensive proteomic and cellular remodeling of the spinal cord—particularly in untrained animals. Prior training buffers both the magnitude and nature of these responses, suggesting that long-term activity impacts spinal cord adaptation.

### Training State Differentially Tunes Glial Transcriptional Programs after Acute Exercise

To address how prior training modulates glial responses to acute exercise, we characterized transcriptional dynamics in spinal cord cell-types 6 and 24h after a single bout of exhaustive exercise. Four glial populations—oligodendrocytes, astrocyte-1, OPCs, and microglia-2— demonstrated extensive DEGs at both timepoints (Figure 4A–C, S5A). At 6h post-exercise, DEGs were predominantly upregulated, with trained mice showing more DEGs. By 24h, DEGs expanded in both groups with a mix of up- and downregulation, representing a biphasic response. Additional glial subtypes, including astrocyte-2 and microglia-1, were also activated with time-dependent patterns, notably with microglia-1 mainly responding at 24h (Figure S5B–C).

**Figure 4.**
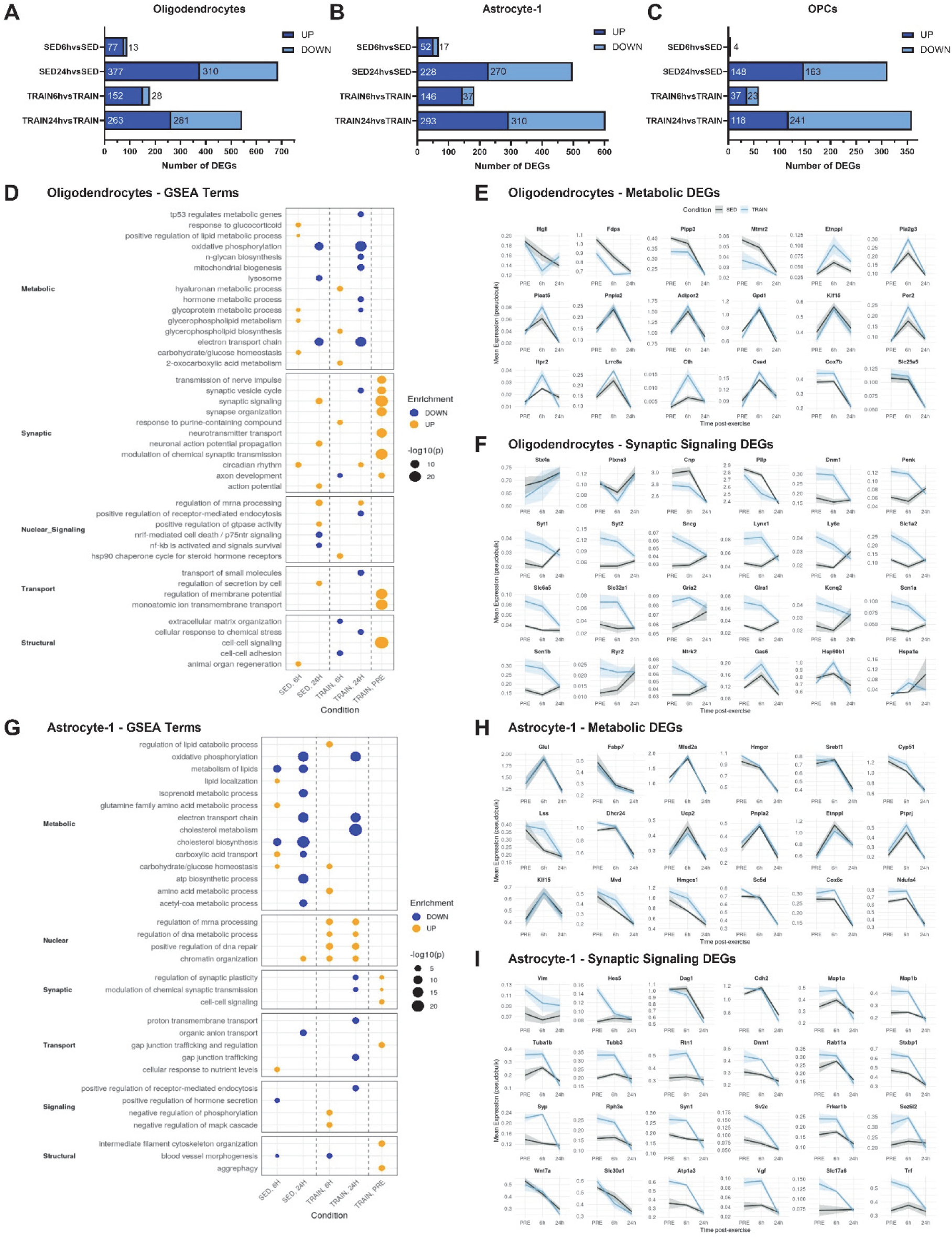
Training Status Tunes Glial Transcriptional Dynamics after Acute Exercise. (A–C) Bar plots show the number of significantly upregulated (dark blue) and downregulated (light blue) differentially expressed genes (DEGs) in oligodendrocytes (A), astrocyte-1 (B), and oligodendrocyte precursor cells (OPCs; C) at 6h and 24h post-acute exercise, comparing sedentary (SED) and trained (TRAIN) mice relative to respective baselines. Genes were considered significant when |log₂FC| > 0.5 and q < 0.01. (D) Gene set enrichment analysis (GSEA) for oligodendrocyte DEGs highlights top enriched functional categories—metabolic, synaptic, structural, signaling, and transport—across timepoints and training states. Circle size reflects statistical significance (–log_10_p), color indicates up- (yellow) or downregulation (blue). (E–F) Line plots depict normalized temporal expression profiles of representative oligodendrocyte DEGs: (E) metabolic genes and (F) synaptic signaling genes across baseline (PRE), 6h, and 24h. Shaded areas represent ± SEM. (G) GSEA for astrocyte-1 DEGs showing dynamically enriched metabolic, synaptic, and signaling pathways by training state and timepoint. (H–I) Line plots depicting expression trajectories for selected metabolic (H) and synaptic signaling (I) genes in astrocyte-1 across conditions. All gene expression data are log-normalized average expression. n = 4 per timepoint and condition.

Overlap analysis confirmed minimal DEG sharing between glial populations (Figure S5D–G), highlighting strong cell-specificity. While oligodendrocytes dominated the baseline transcriptional response to chronic exercise, the acute response strongly engaged astrocyte-1 and OPCs besides oligodendrocytes, particularly in trained mice. This re-weighting suggests that prior training tunes multiple glial types for transient activation following physiological challenge, reflecting a more distributed and responsive glial network.

GSEA demonstrated distinctive functional programs among these glia. Oligodendrocytes were enriched for metabolic and synaptic terms (Figure 4D), with transient upregulation of metabolic genes (*Etnppl, Pla2g3, Plaat5, Itpr2, Cth*) at 6h in both groups, with stronger induction in trained animals, suggesting enhanced metabolic readiness (Figure 4E). Synaptic genes (*Syt1, Syt2*, *Dnm1*, *Gria2*, and *Glra1)* were elevated at baseline in trained animals, remained stable at 6h, and declined at 24h, while sedentary mice showed lower baseline expression and delayed induction at 24h (Figure 4F). These divergent trajectories imply that training pre-establishes a synaptic support state, which undergoes rapid resolution post-exercise, whereas sedentary conditions have a delayed or dampened transcriptional response.

Astrocyte-1 regulated metabolic, synaptic, and signaling genes in a training-dependent manner (Figure 4G), including robust baseline or 6h induction and then suppression at 24h of lipid metabolic (*Fabp7, Pnpla2, Srebf1*) and cholesterol biosynthesis (*Hmgcr, Cyp51, Mvd*) genes (Figure 4H). Mitochondrial genes (*Cox6c, Ndufa4*) were induced at modestly higher levels in trained animals at baseline and 6h (Figure 4H). Genes for synaptic function and cytoskeletal remodeling (*Dnm1, Map1a/b, Rtn1, Tubb3, Rab11a, Syn1, Syp, Sez6l2*), and immediate early/neurotrophic genes (*Vgf, Trf*) were upregulated at baseline in trained mice, then normalized or suppressed by 24h (Figure 4H–I). In sedentary animals, these genes were unchanged or weakly induced at 6h, suggesting that training primes astrocytes for prompt activation and rapid resolution post-exercise.

OPCs exhibited fewer but distinctive changes in metabolic and signaling genes, most notably in trained animals (Figure S5H). At baseline, trained OPCs expressed higher levels of anabolic and cholesterol biosynthesis genes (*Fdps, Msmo1, Hmgcs1, Sqle, Dhcr7/24*), which were downregulated at 6h in TRAIN and at 24h in SED (Figure S5J). Other genes, including those related to glycolysis, redox, chromatin remodeling, and Wnt signaling (*Aldoa, Phka1, Gpd1, Kdm4a, Tcf7l2*), peaked at 6h or 24h in a training-dependent manner. These results indicate that OPCs coordinate anabolic and plasticity programs more efficiently after training.

Microglia-2 exhibited global suppression of classical homeostatic (*P2ry12, Csf1r, Hexb*) and immune (*Tyrobp, C1qc, Ctss, Lamp2*) genes at 24h, regardless of training state (Figure S5I, S5K). A transient upregulation of remodeling- and plasticity-associated genes (*Adora1*, *Tmsb4x*, *Daam2*) at 6h also suggests subtle priming toward a neuroprotective state in response to acute activity.

### Training Modulates the Timing and Nature of Intercellular Communication after Acute Exercise

Applying CellChat L–R analysis across all spinal cord cell-types at baseline, 6h, and 24h post– acute exercise revealed that communication number and strength peaked at 6h and diminished sharply by 24h in both groups (Figure 5A–B), paralleling biphasic transcriptional regulation in glia (Figure 4A–C, S5A-C). Trained mice displayed slightly higher overall inferred interactions at baseline and 6h and lower inferred interactions at 24h.

**Figure 5.**
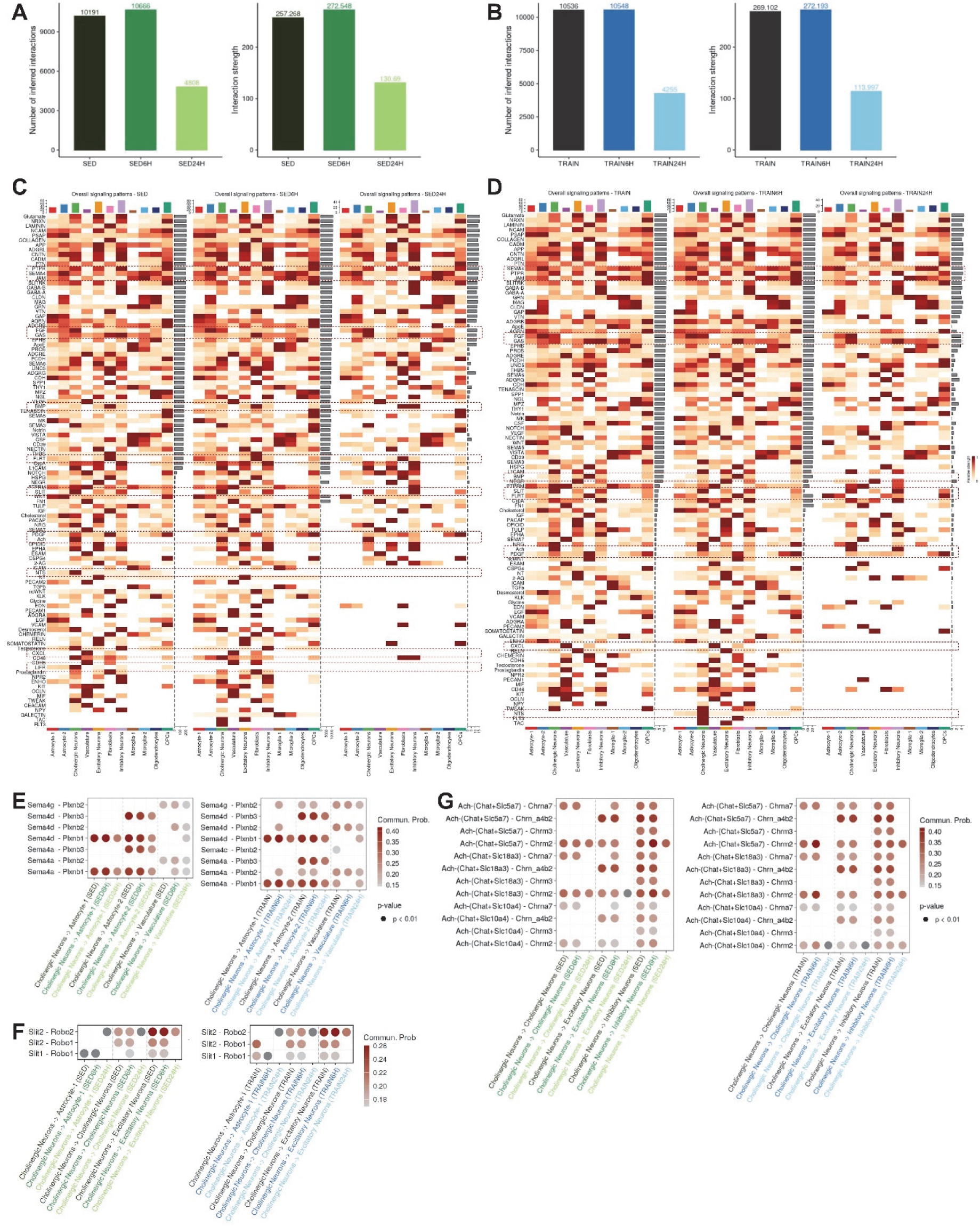
Endurance Training Modulates Dynamic Intercellular Communication Networks in Spinal Cord Following Acute Exercise. (A and B) Bar graphs quantifying the total number (left) and aggregate strength (right) of predicted intercellular communications among spinal cord cell-types inferred by CellChat, across baseline, 6h, and 24h after acute exercise for sedentary (SED, green, A) and trained (TRAIN, blue, B) mice. (C and D) Heatmaps illustrating overall strength of communication for all major signaling pathways for each sender cell-type, highlighting dynamic reorganization of intercellular signaling at baseline, 6h, and 24h in SED (C) and TRAIN (D) conditions. Selected pathways are highlighted with dashed lines. (E–G) Dot plots showing top ligand–receptor interactions mediating Semaphorin (E), Slit (F), and Acetylcholine (G) signaling from cholinergic neurons to glial, vascular, and neuronal targets across conditions. Dot color represents communication probability, all colored dots are statistically significant (p < 0.01), dark grey dots are not significant (p > 0.01).

Pathway-level analysis identified distinct training-dependent remodeling of intercellular communication patterns (Figure 5C–D). In trained animals, positive remodeling was characterized by higher baseline or early activity of acetylcholine, PDGF, FLRT, SLIT, BMP, FGF, PTPR, and SEMA4 signaling, with several of these pathways (e.g., BMP, FGF, PDGF) showing early resolution by 24h. Conversely, sedentary animals preferentially engaged NTS, LIFR, CXCL, JAM, and sustained FGF/PTPR signaling, often peaking at 6h and persisting through 24h, indicative of a prolonged inflammatory/trophic state.

Cholinergic neuron–derived signaling showed marked training-dependent remodeling across multiple pathways (Figure 5E–G, S6). Guidance cues such as Semaphorin and Slit were strongly enhanced in trained animals: Sema4 signaling to astrocytes and vasculature increased particularly at 6h for astrocyte-1, at baseline and 6h for astrocyte-2, and at all timepoints for vascular targets, while Slit signaling rose from cholinergic neurons to both astrocytes and themselves (Figure 5E-F). In both cases, the elevated signaling strength reflected increased expression of receptors in the target populations (astrocytes and cholinergic neurons), rather than increased ligand expression in the sending neurons (Figure S6A–C). This shift suggests that glial cells, often the primary receivers, act as downstream transcriptional responders to neuronal cues. Within neuronal circuits, acetylcholine signaling dynamics further diverged: in trained mice, self-signaling among cholinergic neurons peaked sharply at 6h and resolved by 24h, whereas in sedentary mice it persisted through 24h. By contrast, cholinergic-to-inhibitory signaling was stronger in SED at 6h, while excitatory neuron targeting favored baseline activity in TRAIN (Figure 5G, S6D). Additional trophic pathways also showed accelerated resolution with training. FGF signaling was elevated in trained animals at baseline (toward cholinergic neurons and OPCs) and at 6h (toward excitatory neurons), with corresponding increases in *Fgfr1/3* expression in OPCs (Figure S6E–F). By contrast, sedentary mice showed prolonged FGF activity in multiple, persisting at 24h post-exercise (Figure S6E-G). Similarly, PTPR signaling was enhanced at baseline and 6h in trained mice but diminished by 24h, whereas sedentary animals showed delayed and sustained activity at 24h (Figure S6H). Modest receptor upregulation (*Ptprd, Ptprf, Ptprs*) accompanied these changes (Figure S6F–G). The fine-tuned regulation of PTPR signaling in TRAIN likely reflects balanced OPC proliferation and maturation supporting efficient myelin adaptation.

Together, these results demonstrate that cholinergic neuron-driven signaling to glial cells is one of the most dynamically altered features of the trained spinal cord—both acutely and chronically—and represents a principal upstream driver of adaptive glial remodeling. Receptor-level tuning in glia likely enables the trained spinal cord to remodel rapidly and resolve signaling states more efficiently than in sedentary conditions.

### Glial Populations Fine-Tune Communication Networks in Response to Exercise

Although cholinergic neurons emerged as central hubs for intercellular signaling after exercise (Figure 5, S6), glial populations—astrocyte-1, oligodendrocytes, and microglia-2—also undergo substantial, training-mediated communication remodeling, reflecting a nuanced regulatory role in shaping the spinal cord environment (Figure 6, S7).

**Figure 6.**
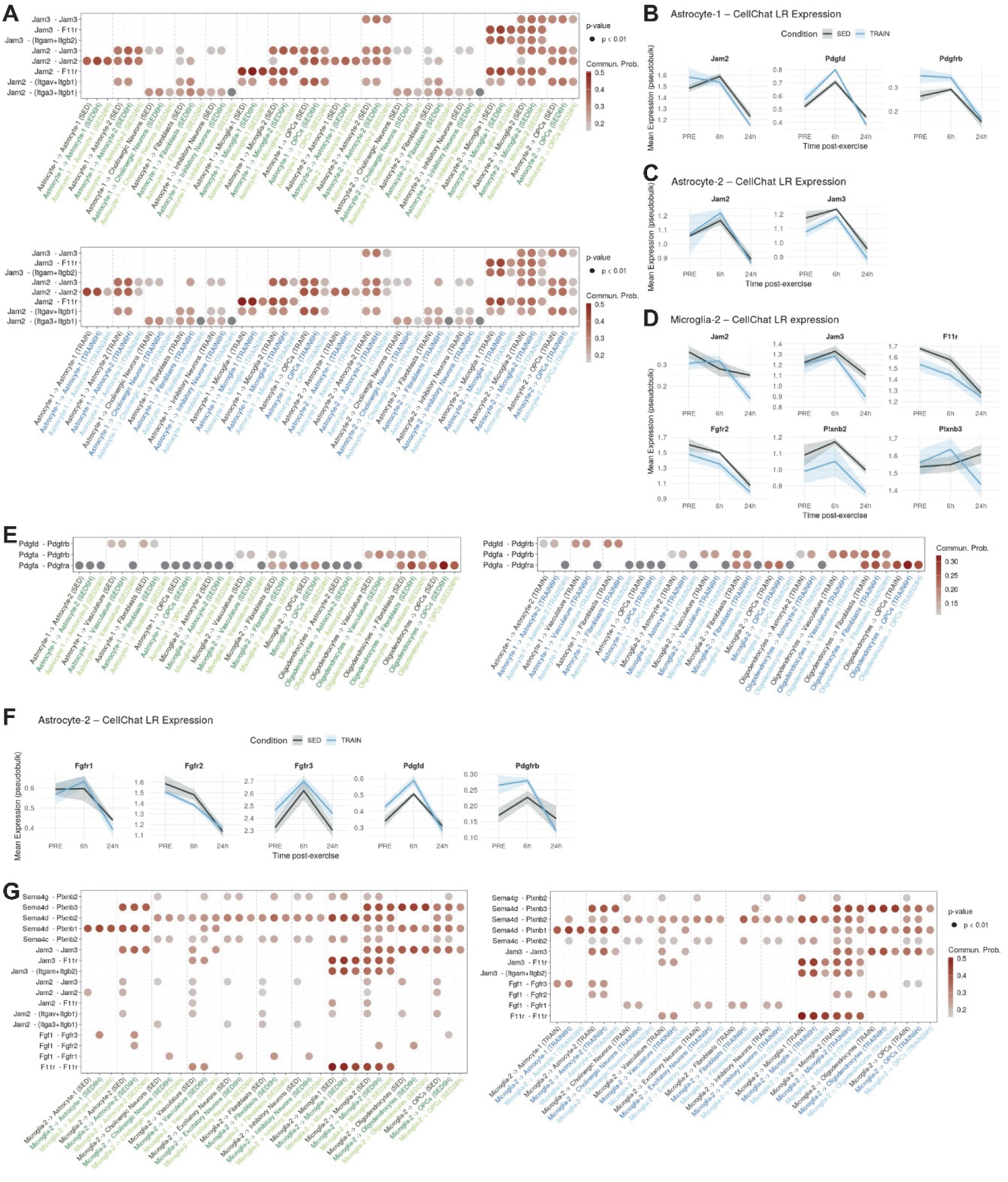
Glial Populations Fine-Tune Intercellular Communication Networks after Exercise in a Training-Dependent Manner. (A) Dot plots display the probability of JAM family ligand–receptor interactions from astrocyte-1 and astrocyte-2 to multiple cell-types by group and timepoint. Top: SED, Bottom: TRAIN. Dot color represents communication probability, all colored dots are statistically significant (p < 0.01), dark grey dots are not significant (p > 0.01). (B–D) Line plots showing mean (±SEM) expression of key JAM (Jam2, Jam3, F11r), PDGF (Pdgfd, Pdgfrb), and FGF (Fgfr2) pathway genes in astrocyte-1 (B), astrocyte-2 (C), and microglia-2 (D), comparing SED and TRAIN animals across PRE, 6h, and 24h post-exercise. (E) Dot plots ligand–receptor probabilities for PDGF signaling from astrocyte-1, microglia-2, and oligodendrocytes to selected target populations across timepoints and condition. Dot color represents communication probability, all colored dots are statistically significant (p < 0.01), dark grey dots are not significant (p > 0.01). (F) Line plots showing mean (±SEM) expression profiles of key FGF (Fgfr1, Fgfr2, Fgfr3) and PDGF (Pdgfd, Pdgfrb) pathway genes in astrocyte-2 in SED and TRAIN conditions. (G) Dot plots of key semaphorin, JAM, and FGF ligand–receptor interactions from microglia-2 to various targets by group and timepoint. Dot color represents communication probability, all colored dots are statistically significant (p < 0.01).

Astrocytes showed pronounced differences in overall signaling between training states. JAM signaling was stronger in sedentary animals, particularly at baseline and 24h, where astrocyte-1 and astrocyte-2 sent enhanced autocrine signals, to microglia-2, and to OPCs (Figure 6A). This was mirrored by higher expression of JAM receptors (*F11r, Jam2, Jam3*) in sedentary astrocytes at these timepoints (Figure 6B–D, S7A). By contrast, semaphorin signaling from astrocytes to other astrocytes and to vascular cells was increased at baseline and 6h post-exercise in trained animals (Figure S7C), mirroring patterns seen in cholinergic neurons (Figure 5E).

Other glial pathways also showed training-specific regulation. PDGF signaling, originating from astrocyte-1, microglia-2, and oligodendrocytes, was enriched in trained mice at baseline and 6h, with higher ligand–receptor expression in astrocytes and OPCs (Figure 6B, 6E–F, S7A–B). These interactions are associated with growth and differentiation programs, suggesting that training primes glia for adaptive remodeling. In contrast, FGF signaling from oligodendrocytes was elevated in sedentary animals at baseline and 6h and persisted at 24h (Figure S7D), whereas trained animals exhibited lower levels at 24h. The broad elevation of oligodendrocyte-derived FGF signaling across multiple spinal cord cell-types in sedentary animals suggests a reliance on prolonged trophic support and maintenance cues, whereas its earlier resolution in trained animals may reflect a shift toward more efficient, differentiation-driven remodeling.

Microglia-2 demonstrated divergent signaling programs, with SED animals showing broadly elevated JAM-mediated inflammatory signaling towards all cell-types at baseline that was resolved following exercise (Figure 6D, G). In contrast, trained microglia showed stronger engagement in growth- and guidance-related pathways—including SEMA4 and FGF—already at baseline; these signals were then transiently downregulated at 6h and 24h post-exercise, respectively. Notably, in SED animals, SEMA4 and FGF signaling emerged only at 6h post-exercise. This pattern indicates that even a single bout of exercise biases microglia away from inflammatory outputs and toward trophic and structural support.

In sum, glial populations act as fine-tuners of the spinal cord environment. While cholinergic neurons appear to provide primary “drive,” glia regulate the timing, precision, and qualitative balance of communication pathways. This dual mechanism allows exercise to orchestrate coordinated, cell-type–specific remodeling—where upstream neuronal cues initiate broad adaptation and glial cells enforce an adaptive balance between inflammation, structure, and trophic support.

## DISCUSSION

Physical activity exerts profound benefits on neural health, yet how the spinal cord, a critical hub for sensorimotor integration, adapts to exercise under healthy conditions has remained largely unexplored. Previous work has focused primarily on exercise in the context of injury, neurodegeneration, aging, or systemic disease, where exercise often mitigates pathology or enhances recovery(7, 9, 10, 13, 14, 16, 17, 19, 50, 51). As examples, two recent studies examined exercise effects in the aging spinal cord using snRNA-seq, demonstrating that aerobic exercise can partially restore youthful cellular signatures and attenuate age-associated microenvironmental decline (Du et al. 2025; Li et al. 2023).

By contrast, the present study provides a multi-omic characterization of how the healthy spinal cord remodels in response to an acute endurance exercise bout in a training state-dependent manner, as well as to chronic training at rest. Using bulk proteomics, fixed tissue snRNA-sequencing, and cell–cell communication inference in the lumbar spinal cord, we uncover transcriptional and signaling programs that engage predominantly in glial populations and are shaped by time post-exercise and training status. Our results highlight glial plasticity and cholinergic neuron–glia crosstalk as central mechanisms through which exercise remodels the spinal cord microenvironment.

### Glia emerge as the primary transcriptional responders

Across analyses, oligodendrocytes, astrocytes, OPCs, and microglia displayed the most extensive transcriptional remodeling, whereas neurons exhibited more modest and heterogeneous responses, placing glia under the spotlight for mediating exercise-induced adaptations (Figure 7A). Functionally, the transcriptional programs engaged across these populations were strikingly convergent—enrichment for metabolic coordination, lipid turnover, synaptic signaling support, and structural remodeling. However, the specific gene programs were largely cell-type specific, underscoring tailored glial contributions (Figure 7A). These findings align with mounting evidence that glia are not merely passive supporters, but active modulators of circuit excitability, synaptic remodeling, and motor performance (52–55).

**Figure 7.**
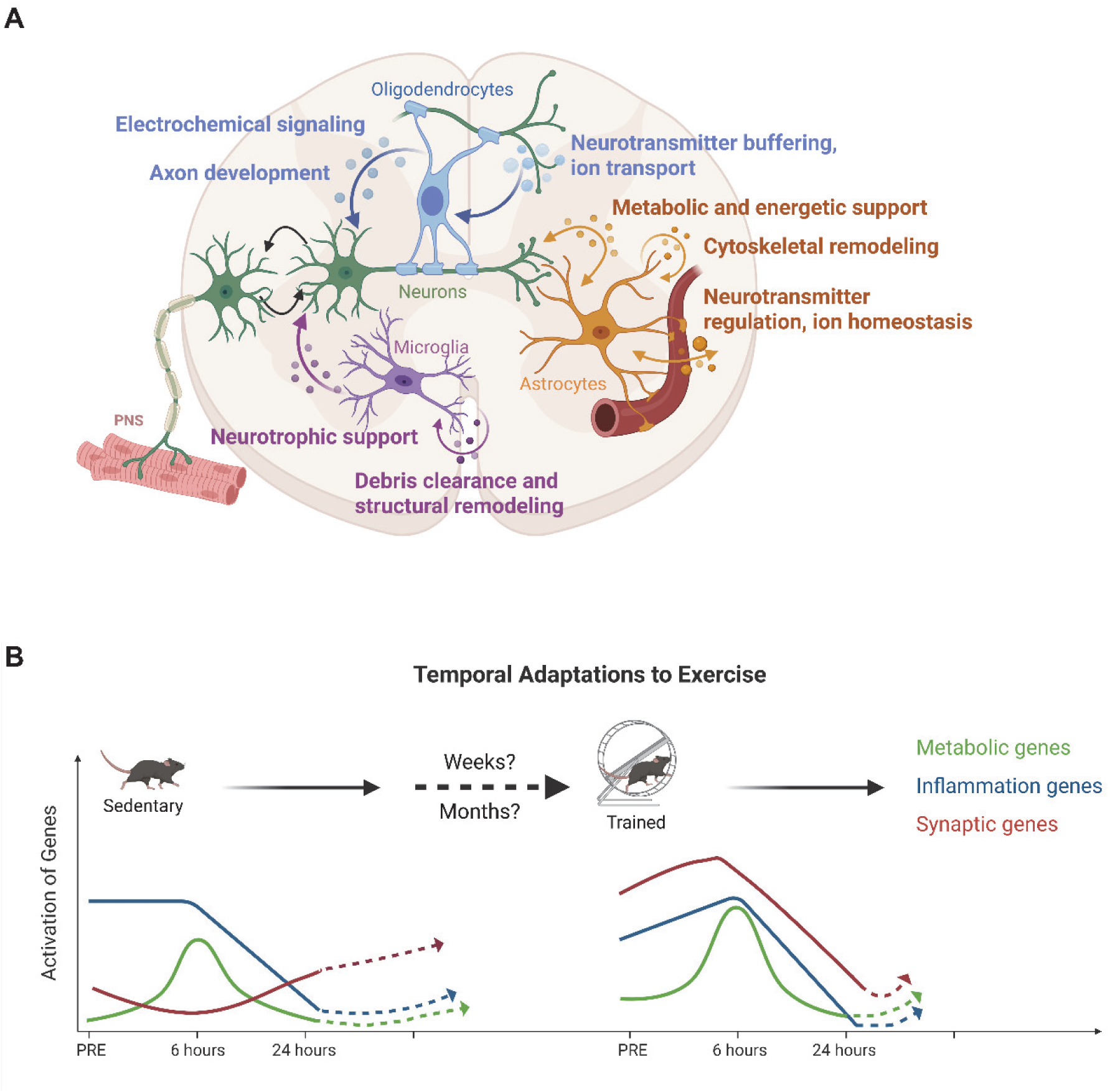
Schematic Summary of Exercise-Induced Transcriptional Programs and Their Temporal Dynamics in the Healthy Lumbar Spinal Cord. (A) Summary of major transcriptional processes modulated by endurance exercise across spinal cord cell types. Arrows indicate upregulated or downregulated pathways identified by snRNA-seq, including metabolic coordination, synaptic support, lipid turnover, cytoskeletal remodeling, and inflammatory or growth-associated signaling. Each cell type is represented by its dominant gene programs and functional themes derived from differential expression and pathway analyses. Abbreviation: PNS, peripheral nervous system. (B) Temporal trajectories of representative gene programs illustrating exercise-induced dynamics in sedentary and trained mice. Curves depict the relative activation of three key functional categories across baseline, 6 h, and 24 h after an acute bout of exercise. Training accelerates the induction and resolution of transcriptional responses, whereas sedentary animals show delayed and more persistent activation. Together, these schematics illustrate the cell-type specificity and temporal logic of spinal cord adaptation to exercise and highlight the influence of prior training on glial and neuronal plasticity. Figures created with BioRender.com

### Training state dictates the temporal dynamics of glial responses

By profiling both steady-state (baseline) adaptations after training at rest and acute responses at 6h and 24h following a single bout of exercise, we distinguished long-lasting remodeling from transient shifts. A clear biphasic pattern emerged: metabolic and signaling genes were induced at 6h and largely resolved by 24h. Crucially, trained animals exhibited earlier induction and faster resolution, whereas sedentary animals showed delayed and more persistent activation. For example, synaptic genes in oligodendrocytes and astrocytes were already elevated at baseline in trained mice, remained stable at 6h, and declined at 24h, while in sedentary animals, they were induced only at 6h for astrocytes and at 24h for oligodendrocytes. These temporal distinctions between trained and sedentary animals are summarized in Figure 7B, which illustrates representative gene programs across key functional categories and highlights how prior training shifts the timing, amplitude, and persistence of transcriptional responses. The training state-dependent re-weighting implies that prior exercise primes glia for rapid but transient responses, thereby reducing the persistence of injury-like or inflammatory phenotypes that characterize sedentary animals. Such dynamics likely underlie the enhanced resilience and adaptive capacity observed in trained spinal cords.

### Neuron–glia communication is a central driver of adaptation

One of the more curious findings was that cholinergic neurons, despite showing few DEGs, emerged as dominant hubs of intercellular signaling. CellChat analysis revealed training-dependent remodeling of cholinergic neuron–glia pathways including Semaphorin, Slit, PDGF, FGF, and PTPR. In trained animals, these signals were engaged at baseline or within 6h and resolved by 24h, whereas in sedentary animals they were absent or persisted at 24h. These pathways converge on processes of axon guidance, synaptic adhesion, vascular remodeling, and myelin adaptation—precisely the themes transcriptionally enriched in glial populations (Figure 7A). The overlap strongly suggests that cholinergic neurons coordinate glial transcriptional programs through paracrine signaling, positioning cholinergic neuron–derived cues as upstream regulators of glial plasticity. This observation parallels recent work implicating motor neurons in shaping oligodendrocyte and astrocyte states via activity-dependent signaling (55–58).

### Microglial remodeling complements astrocyte and oligodendrocyte adaptation

Microglia exhibited a distinct profile characterized by diminished inflammatory JAM signaling at baseline in trained animals alongside early engagement of growth- and guidance-related pathways such as SEMA4 and FGF. This was accompanied by transient induction of remodeling-associated genes such as *Adora1, Tmsb4x, and Daam2*. Functionally, this suggests a shift away from inflammatory activation and toward a more protective, growth-supportive phenotype. This pattern resonates with recent findings that exercise promotes anti-inflammatory microglial states in the brain (59–61) and extends this principle to the spinal cord.

### Mechanistic implications and integration with existing literature

Our findings extend previous work showing that exercise enhances neurogenesis, angiogenesis, and myelin remodeling in the brain (1, 32, 57, 62–64) and injured spinal cord (7, 9–14, 16, 21, 50) by demonstrating that similar principles apply in the healthy spinal cord. The convergence of metabolic and synaptic programs across glial types supports their role as activity sensors and circuit stabilizers and is summarized in the cell-type–specific schematic in Figure 7A. The central contribution of cholinergic neurons highlights a model in which these neurons orchestrate adaptive plasticity indirectly, by modulating glial activity rather than relying on broad neuronal transcriptional changes. At the same time, our work complements recent aging-focused studies that applied snRNA-seq to examine how aerobic exercise influences spinal cord cell states (45, 46). These studies showed that exercise can partially restore youthful transcriptional trajectories in aged animals, but their designs did not address several key features of exercise adaptation in the healthy adult spinal cord. Specifically, neither study captured acute temporal dynamics after an acute exercise bout, assessed inter-individual variability due to pooled sequencing replicates, nor examined lumbar spinal segments directly involved in hindlimb locomotion. Both studies also relied on fresh-tissue dissociation, which can introduce stress-related transcriptional signatures that are minimized by fixed-tissue approaches. By focusing on healthy adult mice, incorporating independent biological replicates, profiling lumbar segments, and resolving transcriptional and signaling programs at 6 h and 24 h after an acute bout of exercise in addition to baseline adaptations, our design captures aspects of exercise-induced plasticity that previous atlases could not detect, e.g. temporal waves of gene expression, inter-individual variability, and region-specific remodeling in lumbar segments innervating hind limbs. These distinctions are reflected in the temporal model presented in Figure 7B, which illustrates how prior training alters the amplitude and resolution of transcriptional responses across major glial populations. Together, these observations position our study as a complementary contribution to the emerging literature, defining the cellular and temporal logic of exercise-induced spinal cord adaptation in a physiological, non-aging context.

### Limitations and future directions

Several limitations should be noted. First, cholinergic neurons were underrepresented in the snRNA-seq dataset, limiting our ability to detect subtle transcriptional changes. Enriching for cholinergic neurons using ChAT-Cre lines, as applied in prior work (41, 44, 65), will be important for future studies. Second, our timepoints (6h, 24h) may have missed very early (0– 4h) or late (>24h) neuronal responses to an acute bout of exercise. Given the timing of peak firing and neuromuscular junction remodeling, expanding the temporal resolution will be critical. Third, although bulk proteomics validated broad proteome-wide changes, cell-type– resolved proteomics or spatial transcriptomics would strengthen the mechanistic links between transcription, translation, and local circuit remodeling. Then, our study was only performed in male mice, using specific endurance exercise and training paradigms. Sex-dependent differences, as well as the effect of other training protocols, for example resistance exercise, will have to be assessed. Finally, while our study documented extensive molecular adaptation, functional validations and assays of spinal circuit excitability or NMJ remodeling will be essential to establish causal links between glial plasticity and motor performance.

### Conclusions

In summary, endurance exercise remodels the healthy spinal cord through coordinated transcriptional plasticity of glia and neuron–glia communication. Training primes oligodendrocytes, astrocytes, and OPCs for rapid metabolic and synaptic responses, accelerates resolution of transcriptional activation, and dampens inflammatory microglial states (Figure 7). Beyond transcription, exercise reshapes communication networks, with cholinergic neurons acting as major signaling hubs that likely drive changes in the transcriptional response of glia. Our findings identify coordinated neuron–glia interactions as fundamental mediators of exercise-induced spinal cord adaptation and provide a framework for understanding how physical activity promotes resilience in neural circuits and benefits motor system function in a healthy state. These findings are important on several levels: first, exercise, in particular when performed to exhaustion, represents a physiological state of prolonged (hyper-) activation of motor systems, most likely contributing to fatigue and therefore exercise cessation (66). Insights on the acute perturbations in untrained and trained mice as well as the persistent chronic adaptations therefore help to understand the molecular and cellular underpinnings of spinal cord plasticity. Second, the pathways and interactions resulting in increased resilience, reduced inflammation, and faster return to baseline in the trained spinal cord, together with findings from other reports describing benefits of physical activity in different contexts of spinal cord pathologies, will help to identify and potentially leverage mechanisms and targets for therapeutic approaches.

## MATERIALS AND METHODS

### Animals

Male C57BL/6J mice (from Janvier Labs) were maintained in the animal facility of the Biozentrum, University of Basel under standard housing conditions (22 °C; 12-h light/dark cycle) with unrestricted access to water and standard chow. Experimental protocols were approved by the Cantonal Veterinary Office Basel-Stadt and complied with Swiss animal welfare regulations. Animals were habituated to the housing environment for at least one week prior to the start of experiments. At 11 weeks of age, mice were randomly divided into sedentary controls or a training group. The training group was given access to running wheels equipped with sensors to continuously log distance, while sedentary mice were provided with plastic housing structures for enrichment. Wheel running was voluntary and monitored for 6 weeks. Body composition was assessed at baseline and 4 weeks after the start of the running wheel period using an EchoMRI-100 system in awake, briefly restrained animals. Body weight was monitored weekly until the end of the experimental duration.

### Exhaustive treadmill test

Endurance capacity was assessed on a motorized treadmill set to a 5° incline at both the beginning and end of the 6-week training phase. The final test was used both to measure adaptation and to provide an acute bout of exercise. Running wheels were removed 48 h before treadmill testing to limit immediate effects of voluntary activity. Mice were acclimatized to the treadmill over two days: day 1 consisted of 5 min each at 0, 5, and 8 m/min, and day 2 of 5 min each at 5, 8, and 12 m/min. On day 3, mice performed the maximal test protocol, starting with 5 min at 5 m/min, followed by 5 min at 8 m/min, and then incremental increases of 2 m/min every 3 min up to 32 m/min. Beyond that, speed was increased by 2 m/min every 10 min until exhaustion. Exhaustion was defined as the inability to resume running despite gentle prodding and brief, mild electrical stimulation (0.5 mA, 200 ms pulses at 1 Hz). Animals were sacrificed either 6 h or 12 h following exhaustion. Mice assigned to the baseline sedentary and trained groups were not exposed to the treadmill to avoid introducing acute transcriptional changes.

### Proteomics sample preparation and analysis

Lumbar spinal cord sections of 0.5 cm were finely minced, suspended in lysis buffer (5% SDS, 10mM TCEP, 0.1 M TEAB) and lysed by sonication using a PIXUL Multi-Sample Sonicator (Active Motif) with Pulse set to 50, PRF to 1, Process Time to 10 min and Burst Rate to 20 Hz. Lysates were incubated for 10 min at 95°C, alkylated in 20 mM iodoacetamide for 30 min at 25°C and proteins digested using S-Trap™ micro spin columns (Protifi) according to the manufacturer’s instructions. Shortly, 12 % phosphoric acid was added to each sample (final concentration of phosphoric acid 1.2%) followed by the addition of S-trap buffer (90% methanol, 100 mM TEAB pH 7.1) at a ratio of 6:1. Samples were mixed by vortexing and loaded onto S-trap columns by centrifugation at 4000 g for 1 min followed by three washes with S-trap buffer. Digestion buffer (50 mM TEAB pH 8.0) containing sequencing-grade modified trypsin (1/25, w/w; Promega, Madison, Wisconsin) was added to the S-trap column and incubate for 1h at 47 °C. Peptides were eluted by the consecutive addition and collection by centrifugation at 4000 g for 1 min of 40 ul digestion buffer, 40 ul of 0.2% formic acid and finally 35 ul 50% acetonitrile, 0.2% formic acid. Samples were dried under vacuum and stored at −20 °C until further use.

Dried peptides were resuspended in 0.1% aqueous formic acid, 0.02% DDM (n-Dodecyl-B-D-maltoside) and subjected to LC–MS/MS analysis using a timsTOF Ultra Mass Spectrometer (Bruker) equipped with a CaptiveSpray nano-electrospray ion source (Bruker) and fitted with a Vanquish Neo (Thermo Fisher Scientific). Peptides were resolved using a RP-HPLC column (100 um × 30 cm) packed in-house with C18 resin (ReproSil Saphir 100 C18, 1.5 um resin; Dr. Maisch GmbH) at a flow rate of 0.4 ul/min and column heater set to 60°C. The following gradient was used for peptide separation: from 2% B to 25% B over 25 min to 35% B over 5 min to 95% B over 0.5 min followed by 1 min at 95% B to 2% B over 0.1 min followed by 5 min at 2% B. Buffer A was 0.1% formic acid in water and buffer B was 80% acetonitrile, 0.1% formic acid in water.

The mass spectrometer was operated in dia-PASEF mode with a cycle time estimate of 0.95 s. MS1 and MS2 scans were acquired over a mass range from 100 to 1700 m/z. A method with 8 dia-PASEF scans separated into 3 ion mobility windows per scan covering a 400-1000 m/z range with 25 Da windows and an ion mobility range from 0.64 to 1.37 Vs cm^2^ was used. Accumulation and ramp time were set to 100 ms, capillary voltage was set to 1600V, dry gas was set to 3 l/min and dry temperature was set to 200 °C. The collision energy was ramped linearly as a function of ion mobility from 59 eV at 1/K0 = 1.6 V s cm^-2^ to 20 eV at 1/K0 = 0.6 V s cm^-2^.

The acquired files were searched using the Spectronaut (Biognosys v19.0) directDIA workflow against a Mus musculus database (consisting of 17085 protein sequences downloaded from Uniprot on 20220222) and 393 commonly observed contaminants. Quantitative fragment ion data (F.Area) was exported from Spectronaut and analyzed using the MSstats R package v.4.13.0. Data was normalised using the default normalisation option “equalizedMedians”, imputed using “AFT model-based imputation” and p-values and q-values for pairwise comparisons were calculated using the limma package.

### Nuclei isolation and FACS

Nuclei were isolated from frozen lumbar spinal cord using a protocol adapted from previous published studies(65, 67). Tissue was homogenized in 1 mL of low-sucrose lysis buffer (320 mM sucrose, 10 mM HEPES pH 8.0, 5 mM CaCl₂, 3 mM magnesium acetate, 0.1 mM EDTA, 1 mM DTT, 0.1% NP-40) with a glass Dounce homogenizer (20 strokes with pestle A followed by 20 strokes with pestle B). Homogenates were incubated on ice for 10 min, filtered through a 40 μm cell strainer, and rinsed with 500 μL lysis buffer. The filtrate was collected in low-binding tubes, briefly spun to recover residual volume, and centrifuged at 500 × g for 5 min at 4 °C. Pellets were resuspended in low-sucrose buffer, spun a second time, and washed in 500 μL of wash buffer (PBS with 1% BSA and 0.2 U/μL SUPERaseIn). For enrichment of intact nuclei, 900 μL of 1.5 M sucrose solution was mixed with 500 μL of the nuclei suspension and layered over 500 μL of 1.5 M sucrose buffer in a low-binding tube. Gradients were centrifuged at 3,000 × g for 40 min at 4 °C (swinging-bucket rotor). Pellets were resuspended in wash buffer and centrifuged for 10 min at 500 × g to remove residual sucrose. The resulting nuclei pellet was fixed for 1 h at room temperature using the 10X Genomics Fixation Kit (#1000497) and quenched according to the manufacturer’s protocol. Nuclei were stained with DAPI and sorted on a BD Aria Fusion cytometer to enrich for DAPI-positive events. A small fraction of sorted nuclei was reserved for imaging quality control, while the remainder was cryopreserved following the 10X Genomics Flex protocol (CG000478 | Rev C) and stored at −80 °C until library preparation.

### Single-nucleus library preparation

Single-nucleus RNA-seq was performed with the Chromium Next GEM Fixed RNA kit (10X Genomics; User Guide Rev D, CG000522). Nuclei were counted using Acridine Orange/Propidium Iodide staining on a LUNA-FL fluorescence counter. After probe hybridization, samples were pooled to ensure equal representation (pool-and-wash workflow). Each run included 16 multiplexed samples, with a minimum of 80’000 nuclei loaded per sample. Library quality was confirmed using a Bioanalyzer, and sequencing was carried out on an Illumina NovaSeq6000 platform targeting approximately 15’000 reads per nucleus.

### Single-nucleus RNA-seq data processing and analysis Preprocessing, quality control, data integration, and clustering

Feature-barcode matrices were generated with the Cell Ranger pipeline (10X Genomics, v7.1.0). Ambient RNA contamination was removed using CellBender (v0.3.0)(68). Count matrices were imported into Seurat v5 (69) for downstream analysis. Nuclei with fewer than 500 or more than 8,000 detected genes, >25,000 UMI counts, or mitochondrial read content above 1% were excluded. Putative doublets were identified using DoubletFinder, and clusters enriched for doublet scores were removed. Data were normalized with *SCTransform*, regressing out sequencing depth and mitochondrial percentage. Integration across replicates and experimental conditions was performed using Seurat’s IntegrateLayers with batch correction via Harmony. Dimensionality reduction was conducted with PCA followed by UMAP, and clusters were identified using the Louvain algorithm at a resolution of 0.5.

Cell-types were annotated based on canonical marker expression for excitatory neurons (*Slc17a6, Slc17a8*), inhibitory neurons (*Gad1, Gad2, Slc6a5*), cholinergic neurons (*Chat, Slc18a3, Slc5a7*), astrocytes (*Gfap, Aldh1l1, Aqp4*), oligodendrocytes (*Mbp, Plp1, Mag*), oligodendrocyte precursor cells (*Pdgfra, Cspg4*), microglia (*P2ry12, Tmem119*), vascular cells (*Pecam1, Flt1*), fibroblasts (*Pdgfrb, Vtn*), and pericytes (*Rgs5*). Contaminant clusters were excluded, and closely related clusters were merged into major cell classes.

### Pseudobulk differential expression analysis

For each biological replicate, raw UMI counts were aggregated by cell-type and condition to construct pseudobulk count matrices. DESeq2 (70) was used for differential expression testing across training-state and time course comparisons:

- Training effect: TRAIN vs SED
- Acute sedentary responses: SED6H vs SED, SED24H vs SED
- Acute trained responses: TRAIN6H vs TRAIN, TRAIN24H vs TRAIN.

Lowly expressed genes (row sum < 10 across all samples) were excluded. DESeq2 models included condition and time as factors (∼ condition + time + condition:time). Genes were considered significantly regulated if they had an adjusted p-value < 0.01, absolute log₂ fold change ≥ 0.5, and baseMean ≥ 10. Log₂ fold-change shrinkage was applied using the “*ashr”* method to stabilize estimates.

### Gene set enrichment analysis

Ranked DEG lists were analyzed with *clusterProfiler* using MSigDB gene sets (Hallmark, Reactome, WikiPathways, and GO: Biological Process). Enrichment was performed separately for up- and downregulated genes, and results were visualized as dot plots, heatmaps, and UpSet plots to highlight pathway overlap across cell-types and conditions. Complementary ontology-based enrichment was conducted using the Metascape platform (metascape.org)(71) with default parameters. Where appropriate, custom background gene lists were supplied to account for cell-type–specific expression repertoires. Metascape clusters enriched terms based on shared gene membership and hierarchical clustering, and representative terms from each cluster were selected for visualization.

### Line plot visualization of expression dynamics

To visualize temporal expression changes of selected genes, pseudobulk counts were aggregated per replicate, condition, and timepoint (PRE, 6 h, 24 h). Expression values were converted to counts per million (CPM), log-transformed, and summarized as the mean ± standard error across replicates. Line plots were generated with *ggplot2*, displaying mean trajectories per gene and condition with shaded ribbons representing standard error. Genes were shown in faceted panels with free y-axis scaling, and consistent color schemes and formatting were applied to produce publication-quality figures.

### Cell–cell communication analysis

Intercellular communication analysis was performed using the CellChat R package (version 2.1.2)(49), following the standard comparative workflow. Processed Seurat objects were subset by condition and time point (SED, SED6H, SED24H, TRAIN, TRAIN6H, TRAIN24H) and converted into CellChat objects using the *createCellChat* function, grouping cells by curated major clusters. The CellChat mouse ligand–receptor database was applied. For each subset, we identified overexpressed genes and interactions, inferred communication probabilities, filtered out low-support interactions (<10 cells per partner), and aggregated pathway-level networks.

To enable temporal and training-state comparisons, CellChat objects were merged within sedentary and trained groups and across matched time points. Comparative analyses included quantification of the number and strength of interactions, ranking of pathways by contribution, and visualization of signaling roles using bubble plots, chord diagrams, and role heatmaps. Pathway similarity networks were computed using both functional and structural similarity, followed by UMAP embedding and network clustering to identify groups of related signaling pathways. Network centrality analysis was applied to determine dominant sending and receiving populations across conditions. Direct pairwise comparisons between matched sedentary and trained time points (e.g., SED24H vs TRAIN24H) were also performed to highlight training-dependent differences in signaling patterns.

### Statistics

Differential gene expression was performed using DESeq2 in a pseudobulk framework, with Wald tests for pairwise comparisons and Benjamini–Hochberg correction for multiple testing. Genes with adjusted p-value < 0.01, baseMean ≥ 10, and absolute log₂ fold change ≥ 0.5 were considered significantly regulated. For interaction terms, DESeq2’s likelihood ratio test (LRT) was used to assess the effect of training state on time-dependent responses.

For non-sequencing data (e.g., body composition, behavioral readouts), comparisons between two groups were performed using unpaired Student’s t-tests when normality was confirmed (Shapiro–Wilk test) or Mann–Whitney U tests otherwise. Multi-factorial comparisons (e.g., endurance performance, cell-type proportions, regulon activity) were analyzed using two-way ANOVA followed by Tukey’s post hoc test. Unless otherwise specified, n = 4 mice per group.

## Author Contributions

Conceptualization: S.M. and C.H. Methodology: S.M., S.D., D.R, and C.H. Investigation: S.M., S.D., G.S., and S.A.S. Formal analysis and visualization: S.M. and D.R. Writing: S.M. and C.H. Funding acquisition: C.H. All authors have read and approved the final version of this manuscript and agree to be accountable for all aspects of the work in ensuring that questions related to the accuracy or integrity of any part of the work are appropriately investigated and resolved. All persons designated as authors qualify for authorship, and all those who qualify for authorship are listed.

## Data Availability

Proteomic data have been deposited to the ProteomeXchange Consortium (https://www.proteomexchange.org/) via the MassIVE partner repository (https://massive.ucsd.edu/) with MassIVE data set identifier MSV000099207 and ProteomeXchange identifier PXD068447. The transcriptomics data have been deposited to the Gene Expression Omnibus (GEO) and are available under the accession number XXX.

## Declaration of generative AI and AI-assisted technologies in the writing process

During the preparation of this manuscript, the authors used **ChatGPT** (o4, 5) and **Perplexity** to assist with grammar checking, sentence structure optimization, and improving clarity. These tools were used to refine the language and presentation of ideas. However, the intellectual content, data interpretation, and conclusions remain entirely the authors’ own. The final manuscript was thoroughly reviewed and edited by the authors, who take full responsibility for its content.

## Acknowledgements

We thank the FACS Core Facility of the Biozentrum, especially Stella Stefanova, for assistance with flow sorting and all animal caretakers from the animal facility at the Biozentrum for their help. We thank the Quantitative Genomics Facility of the University of Basel, in particular Phillippe Demougin, for preparing cDNA libraries. Calculations were performed at sciCORE, the scientific computing center at University of Basel, with support by the SIB - Swiss Institute of Bioinformatics.

## SUPPLEMENTARY DATA

**Supplementary Figure S1.**
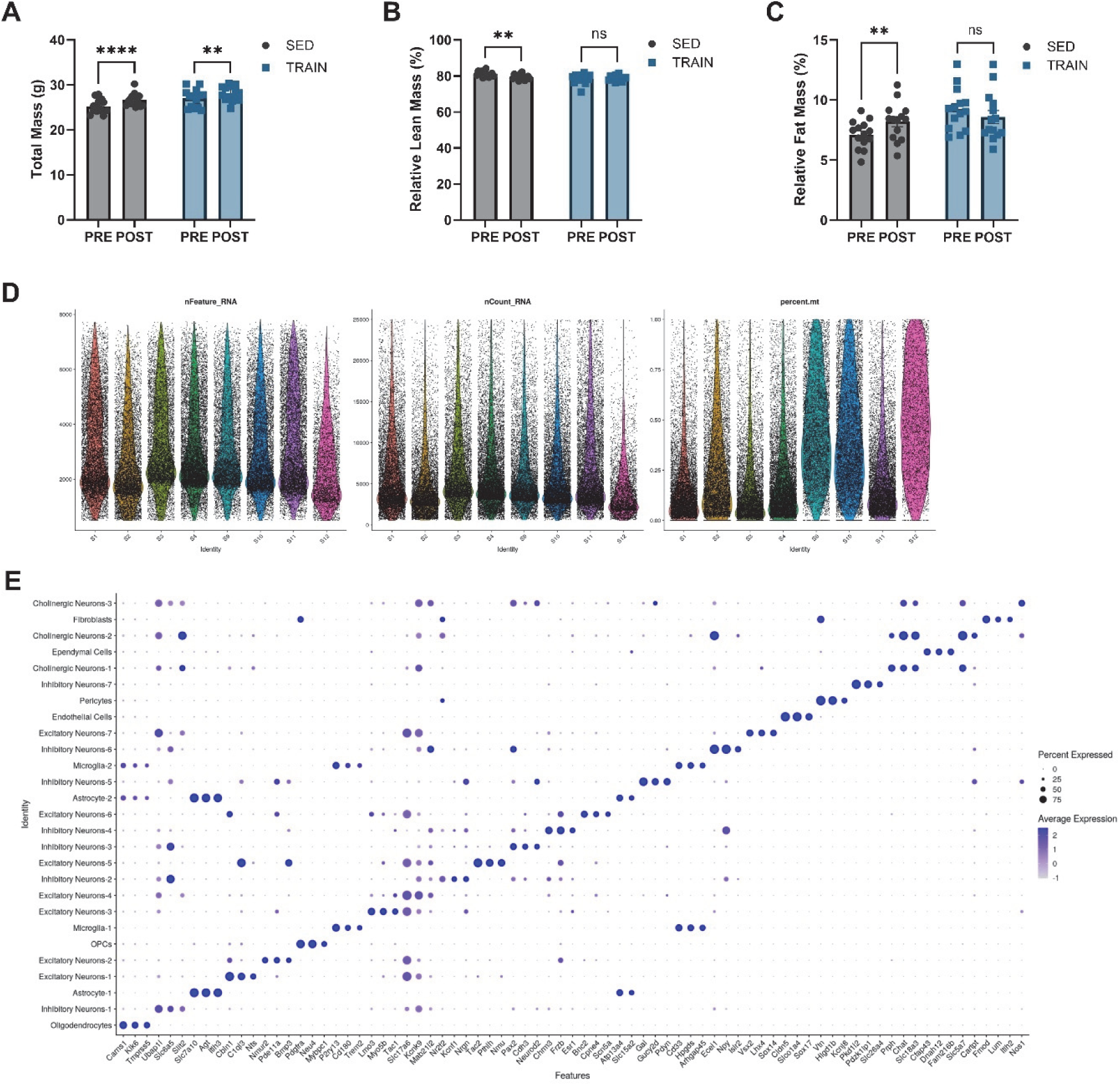
Endurance Exercise Selectively Preserves Lean Mass and Modifies Spinal Cord Cell-Type Landscape. (A–C) Longitudinal assessment of whole-body composition (A, total mass; B, relative lean mass; C, relative fat mass) in SED and TRAIN groups pre- and post-intervention. Data represent mean ± SEM; individual biological replicates shown. **p < 0.01; ****p < 0.0001; ns, not significant; two-way ANOVA with multiple comparisons (Fisher’s LSD test). (D) Violin plots displaying snRNA-seq quality metrics across identified spinal cord cell types: number of genes (nFeature_RNA), unique molecular identifiers (nCount_RNA), and mitochondrial proportion (percent.mt). (E) Dot plot summarizing top marker gene expression of each identified celltype and subcluster in the spinal cord; size of dots indicates percent of cells expressing each marker, color gradient reflects average normalized expression.

**Supplementary Figure S2.**
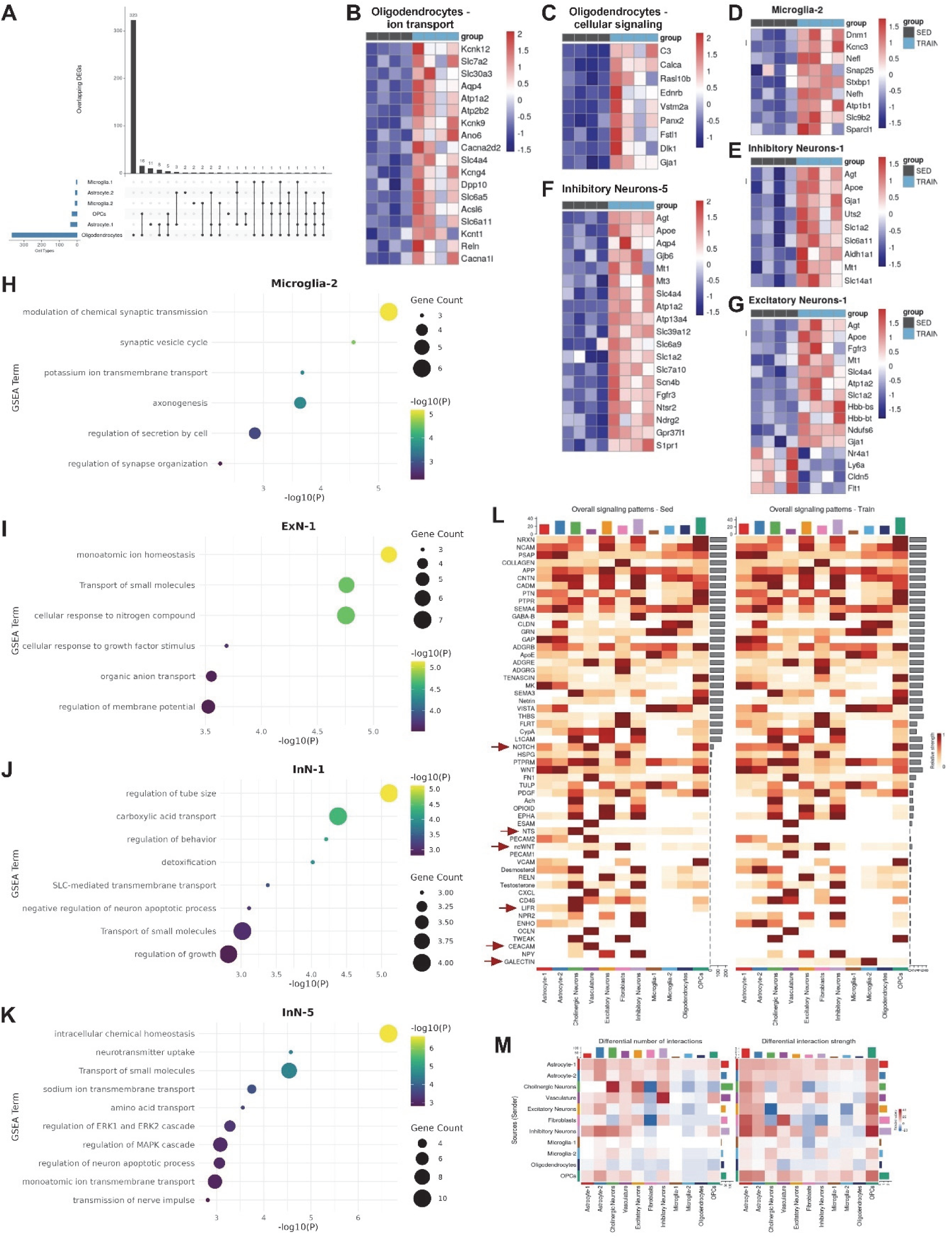
Distinct and Overlapping Transcriptional Programs Across Spinal Cord Cell Types Following Endurance Training. (A) UpSet plot demonstrating limited overlap among differentially expressed genes upregulated in main glial cell types following endurance training. All DEGs analyzed were filtered for |log₂FC| > 0.5 and q < 0.01. (B–C) Heatmaps show relative expression of upregulated genes involved in ion transport (B) and cellular signaling (C) in oligodendrocytes, comparing sedentary (SED) and trained (TRAIN) groups; upregulation is indicated in red and downregulation in blue. (D–G) Heatmaps display top DEG expression profiles in microglia-2 (D), Inhibitory Neurons-1 (E), Inhibitory Neurons-5 (F), and Excitatory Neurons-1 (G). (H–K) Dot plots present selected gene set enrichment analysis (GSEA) results for Microglia-2 (H), ExN-1 (I), InhN-1 (J), and InhN-5 (K), indicating significantly enriched pathways, gene counts, and statistical significance (represented by dot size and color). (L) Heatmaps summarizing overall outgoing signaling pathway patterns and major differences between SED and TRAIN spinal cords, with selected pathways highlighted (red arrows). (M) Interaction heatmaps illustrating differences in the number (left) and strength (right) of intercellular communications between SED and TRAIN, grouped by sending and receiving cell types.

**Supplementary Figure S3.**
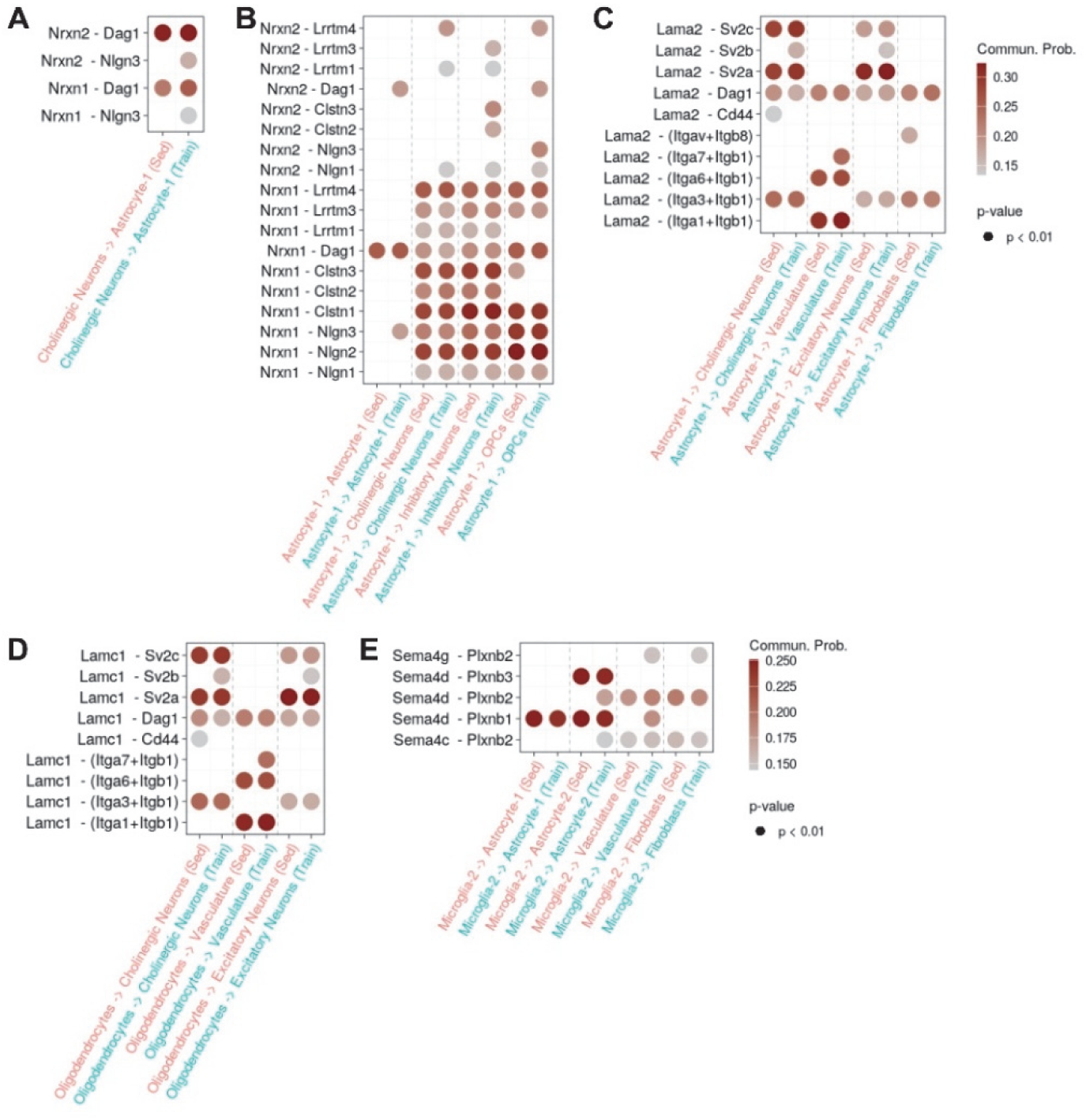
CellChat Analysis Reveals Exercise-Driven Remodeling of Spinal Cord Cell–Cell Signaling Pathways. (A–E) Dot plots display the top CellChat-inferred ligand–receptor interactions mediating signaling from cholinergic neurons (A), astrocytes (B–C), oligodendrocytes (D), and microglia-2 (E) to diverse recipient cell populations in sedentary (SED) and trained (TRAIN) spinal cords. Notable exercise-responsive signaling axes highlighted include Neurexin (Nrxn1/2), Laminin (Lama2, Lamc1), and Semaphorin (Sema4) pathways. Dot color represents communication probability, and dot size indicates statistical significance (all plotted interactions: p < 0.01)

**Supplementary Figure S4.**
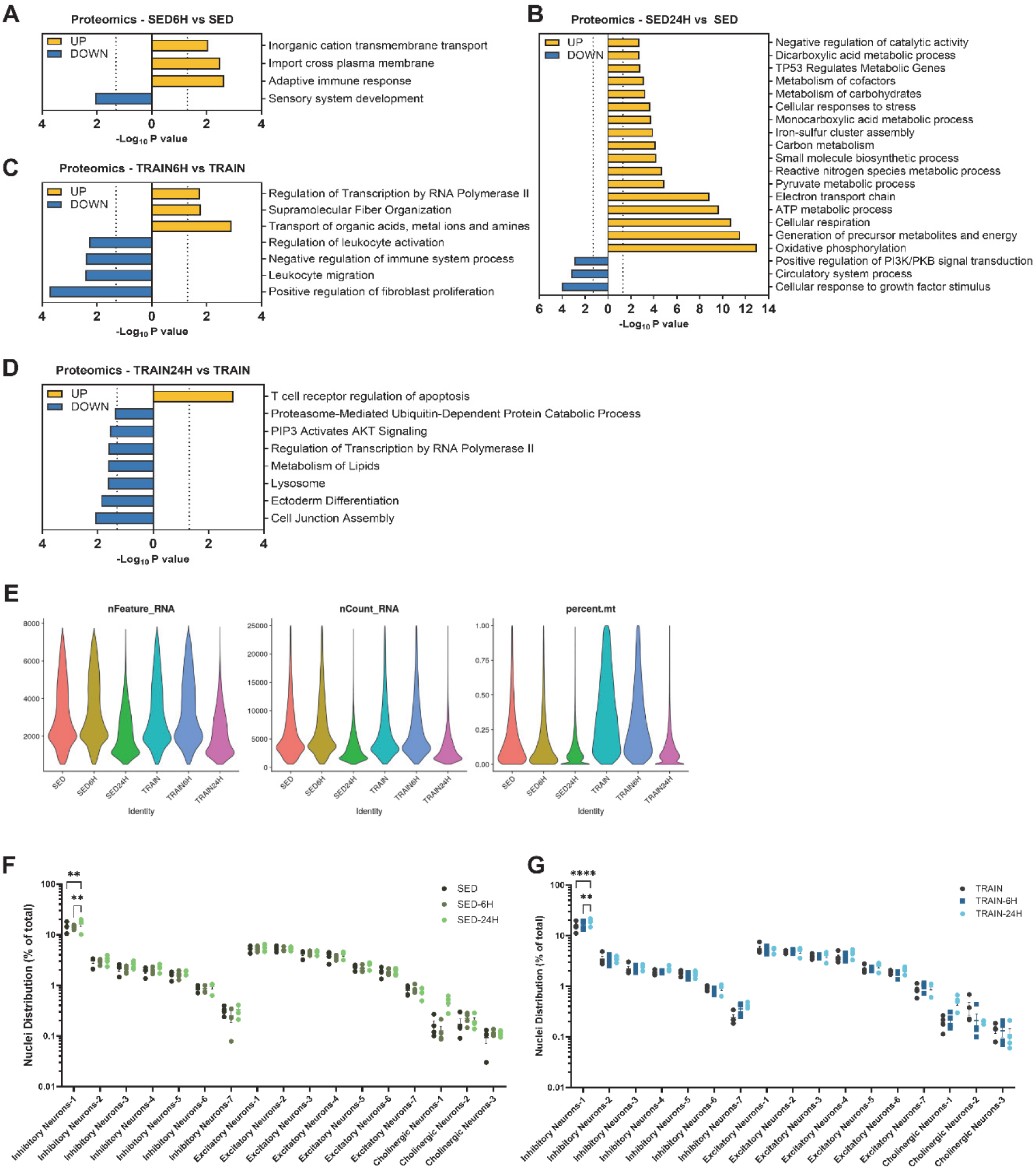
Acute Exercise Modulates Molecular Pathways and Cell-Type Proportions in a Training- and Time-Dependent Manner. (A–D) Gene Ontology (GO) enrichment analysis of significantly up- and down-regulated proteins following acute exercise as determined by bulk proteomics: (A) SED at 6h post-exercise, (B) SED at 24h, (C) TRAIN at 6h, (D) TRAIN at 24h. (E) Violin plots depict key quality control metrics from integrated single-nucleus RNA-seq, including total detected genes (nFeature_RNA), UMI count (nCount_RNA), and percentage mitochondrial reads (percent.mt) across groups and conditions. (F–G) Quantification of neuronal cell-type proportions across conditions: (F) SED baseline, 6h, 24h; (G) TRAIN baseline, 6h, 24h. Proportions plotted as percent of total nuclei (mean ± SEM, n = 4/group). *p < 0.05, **p < 0.01, ****p < 0.0001, ns = not significant (two-way ANOVA with Tukey’s multiple comparisons test).

**Supplementary Figure S5.**
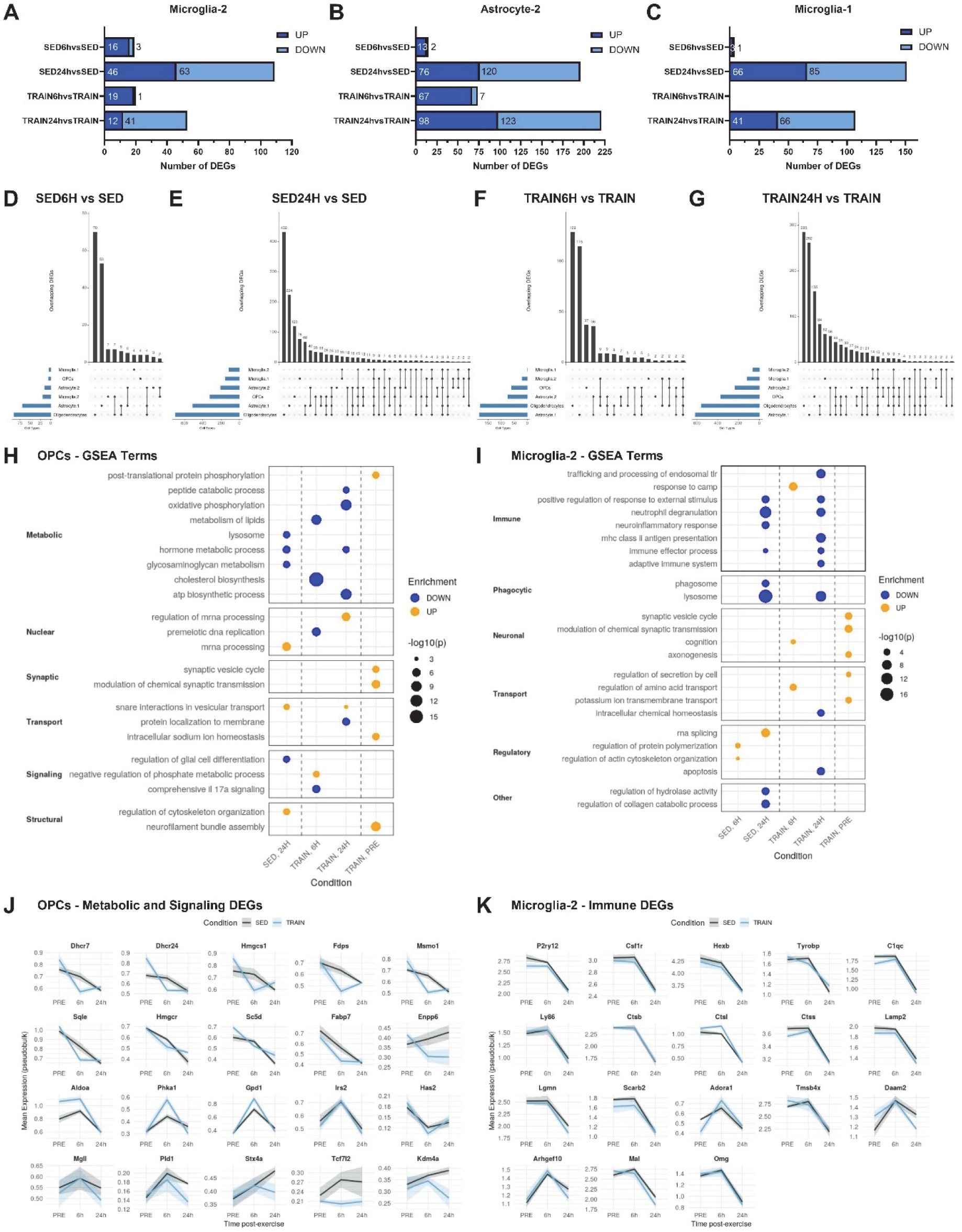
Training Modulates Temporal and Cell-Type–Specific Glial Transcriptional Programs after Acute Exercise. (A–C) Bar plots displaying the number of significantly upregulated (dark blue) and downregulated (light blue) differentially expressed genes (DEGs) in microglia-2 (A), astrocyte-2 (B), and microglia-1 (C) at 6h and 24h post-acute exercise in sedentary (SED) and trained (TRAIN) groups. (D–G) UpSet plots showing overlap of DEGs between main glial subtypes for SED-6h vs SED (D), SED-24h vs SED (E), TRAIN-6h vs TRAIN (F), and TRAIN-24h vs TRAIN (G), highlighting generally limited overlap. (H) Gene set enrichment analysis (GSEA) for OPC DEGs highlights top enriched functional categories—metabolic, nuclear, synaptic, transport, signaling, and structural—across timepoints and training states. Circle size reflects statistical significance (–log10 p), color indicates up- (yellow) or downregulation (blue). (I) GSEA of microglia-2 DEGs for immune, phagocytic, and regulatory categories across timepoints and training states. Circle size reflects statistical significance (–log10 p), color indicates up- (yellow) or downregulation (blue). (J–K) Line plots showing normalized expression trajectories (mean ± SEM) for selected DEGs: representative metabolic and signaling genes in OPCs (J), and immune-related genes in microglia-2 (K), for SED vs. TRAIN animals across baseline, 6h, and 24h post-exercise. All gene expression data are log-normalized average expression. n = 4 per timepoint and condition.

**Supplementary Figure S6.**
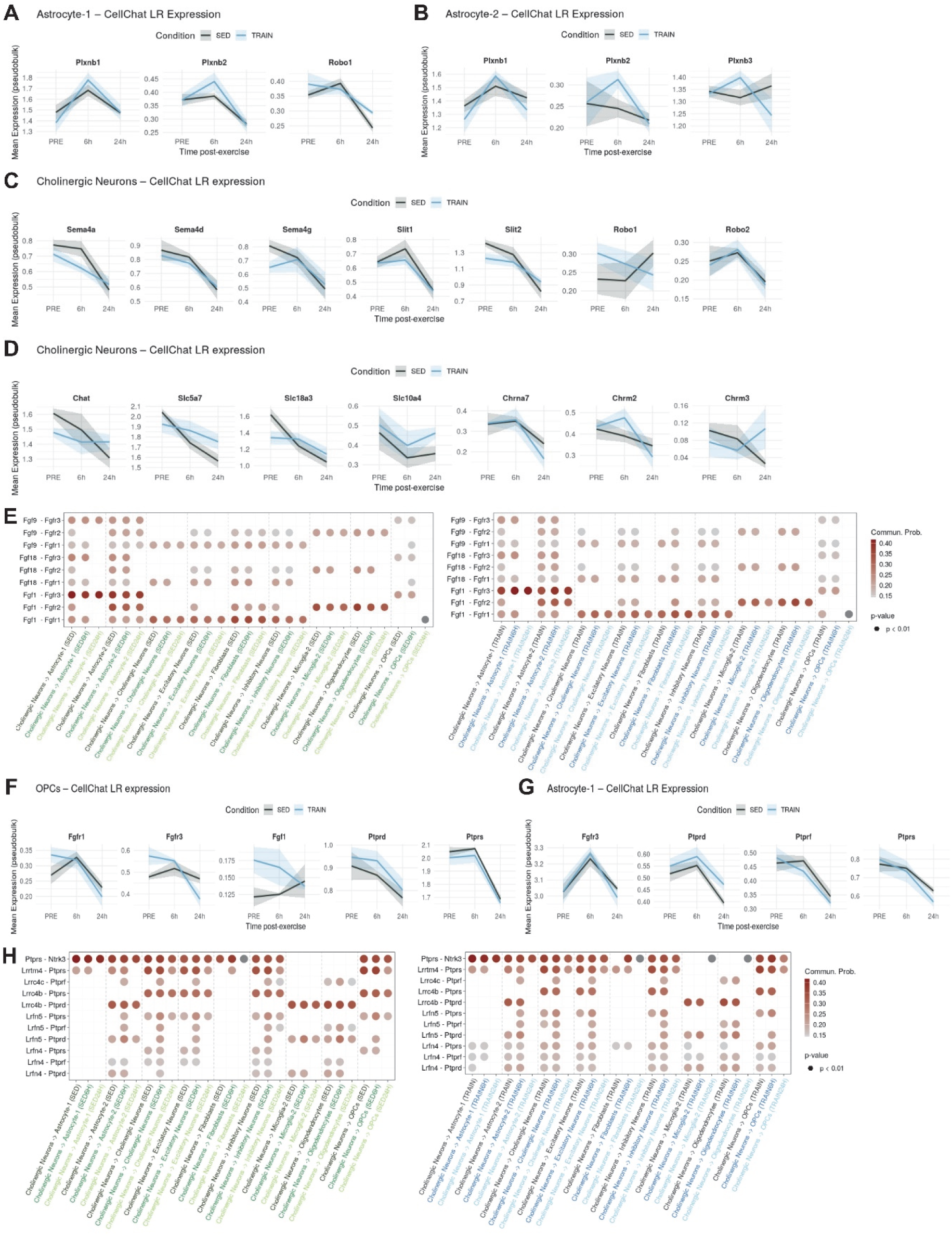
Receptor-Level Regulation Underlies Rapid Remodeling of Glial and Neural Communication After Training. (A and B) Line plots displaying mean (±SEM) expression of Semaphorin and Slit pathway receptors in Astrocyte-1 (A) and Astrocyte-2 (B) across experimental groups and timepoints. (C and D) Line plots displaying mean (±SEM) expression of Semaphorin, Slit, and cholinergic ligand and receptor genes in cholinergic neurons under SED and TRAIN conditions. (E and H) Dot plots indicating CellChat-inferred probabilities for FGF (E), and PTPR (H) ligand– receptor interactions from cholinergic neurons to all major neural and glial cell types by group and timepoint. Dot color represents communication probability, all colored dots are statistically significant (p < 0.01), dark grey dots are not significant (p > 0.01). (F and G) Line plots for OPCs (F) and Astrocyte-1 (G) showing mean expression (±SEM) of key FGF receptors (Fgfr1, Fgfr3) and PTPR family members (Ptprd, Ptprf, Ptprs) over time in SED and TRAIN.

**Supplementary Figure S7.**
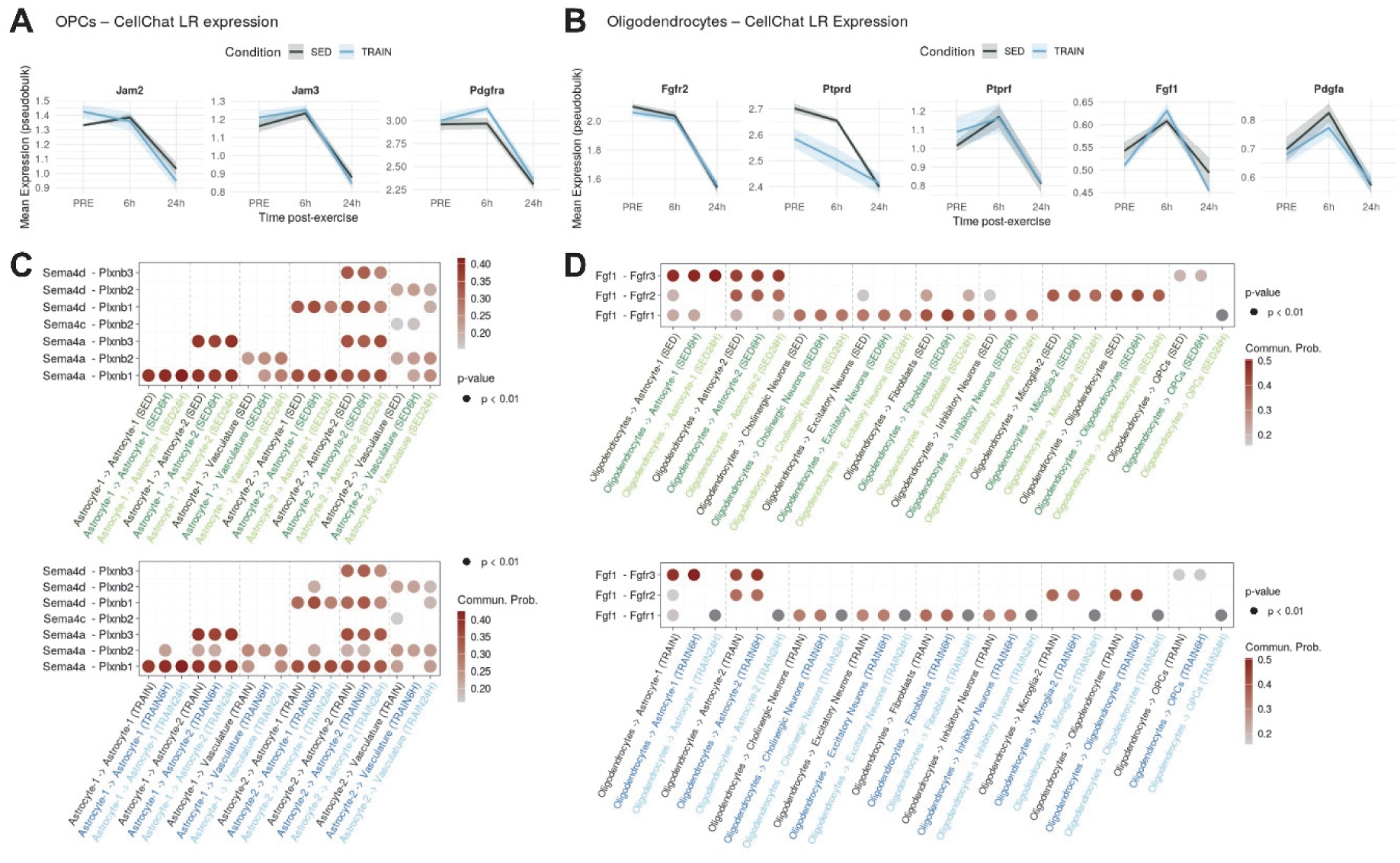
Training State Selectively Regulates Glial and Microglial Communication with the Spinal Cord Niche. (A–B) Line plots showing mean (±SEM) expression profiles of select CellChat ligand/receptor genes in OPCs (A) and oligodendrocytes (B) under SED and TRAIN conditions, illustrating differences in signaling dynamics. (C–D) Dot plots of Semaphorin ligand–receptor interactions sent from astrocyte-1 (C) and FGF ligand–receptor interactions from oligodendrocytes (D) to selected cell types. Top: SED, Bottom: TRAIN. Dot color represents communication probability, all colored dots are statistically significant (p < 0.01), dark grey dots are not significant (p > 0.01).

## REFERENCES

1. van Praag H, Kempermann G, Gage FH. Running increases cell proliferation and neurogenesis in the adult mouse dentate gyrus. Nat Neurosci. 1999;2(3):266–70.

2. Voss MW, Nagamatsu LS, Liu-Ambrose T, Kramer AF. Exercise, brain, and cognition across the life span. J Appl Physiol (1985). 2011;111(5):1505–13.

3. Cotman CW, Berchtold NC. Exercise: a behavioral intervention to enhance brain health and plasticity. Trends Neurosci. 2002;25(6):295–301.

4. El-Sayes J, Harasym D, Turco CV, Locke MB, Nelson AJ. Exercise-Induced Neuroplasticity: A Mechanistic Model and Prospects for Promoting Plasticity. Neuroscientist. 2019;25(1):65–85.

5. Hamilton GF, Rhodes JS. Exercise Regulation of Cognitive Function and Neuroplasticity in the Healthy and Diseased Brain. Prog Mol Biol Transl Sci. 2015;135:381–406.

6. Rossignol S, Dubuc R, Gossard JP. Dynamic sensorimotor interactions in locomotion. Physiol Rev. 2006;86(1):89–154.

7. Bilchak JN, Caron G, Cote MP. Exercise-Induced Plasticity in Signaling Pathways Involved in Motor Recovery after Spinal Cord Injury. Int J Mol Sci. 2021;22(9).

8. Kiss Bimbova K, Bacova M, Kisucka A, Galik J, Zavacky P, Lukacova N. Activation of Three Major Signaling Pathways After Endurance Training and Spinal Cord Injury. Mol Neurobiol. 2022;59(2):950–67.

9. Cobianchi S, Arbat-Plana A, Lopez-Alvarez VM, Navarro X. Neuroprotective Effects of Exercise Treatments After Injury: The Dual Role of Neurotrophic Factors. Curr Neuropharmacol. 2017;15(4):495–518.

10. Cote MP, Azzam GA, Lemay MA, Zhukareva V, Houle JD. Activity-dependent increase in neurotrophic factors is associated with an enhanced modulation of spinal reflexes after spinal cord injury. J Neurotrauma. 2011;28(2):299–309.

11. Flynn JR, Dunn LR, Galea MP, Callister R, Callister RJ, Rank MM. Exercise training after spinal cord injury selectively alters synaptic properties in neurons in adult mouse spinal cord. J Neurotrauma. 2013;30(10):891–6.

12. Jensen SK, Michaels NJ, Ilyntskyy S, Keough MB, Kovalchuk O, Yong VW. Multimodal Enhancement of Remyelination by Exercise with a Pivotal Role for Oligodendroglial PGC1alpha. Cell Rep. 2018;24(12):3167–79.

13. Su H, Luo H, Wang Y, Zhao Q, Zhang Q, Zhu Y, et al. Myelin repair of spinal cord injury in adult mice induced by treadmill training upregulated peroxisome proliferator-activated receptor gamma coactivator 1 alpha. Glia. 2024;72(3):607–24.

14. Engesser-Cesar C, Anderson AJ, Basso DM, Edgerton VR, Cotman CW. Voluntary wheel running improves recovery from a moderate spinal cord injury. J Neurotrauma. 2005;22(1):157–71.

15. Lee DH, Cao D, Moon Y, Chen C, Liu NK, Xu XM, et al. Enhancement of motor functional recovery in thoracic spinal cord injury: voluntary wheel running versus forced treadmill exercise. Neural Regen Res. 2025;20(3):836–44.

16. Lozinski BM, de Almeida LGN, Silva C, Dong Y, Brown D, Chopra S, et al. Exercise rapidly alters proteomes in mice following spinal cord demyelination. Sci Rep. 2021;11(1):7239.

17. Loy K, Schmalz A, Hoche T, Jacobi A, Kreutzfeldt M, Merkler D, et al. Enhanced Voluntary Exercise Improves Functional Recovery following Spinal Cord Injury by Impacting the Local Neuroglial Injury Response and Supporting the Rewiring of Supraspinal Circuits. J Neurotrauma. 2018;35(24):2904–15.

18. Dai Y, Cheng Y, Ge R, Chen K, Yang L. Exercise-induced adaptation of neurons in the vertebrate locomotor system. J Sport Health Sci. 2024;13(2):160–71.

19. Grau JW, Huie JR, Lee KH, Hoy KC, Huang YJ, Turtle JD, et al. Metaplasticity and behavior: how training and inflammation affect plastic potential within the spinal cord and recovery after injury. Front Neural Circuits. 2014;8:100.

20. Chang W, Pedroni A, Bertuzzi M, Kizil C, Simon A, Ampatzis K. Locomotion dependent neuron-glia interactions control neurogenesis and regeneration in the adult zebrafish spinal cord. Nat Commun. 2021;12(1):4857.

21. Krityakiarana W, Espinosa-Jeffrey A, Ghiani CA, Zhao PM, Topaldjikian N, Gomez-Pinilla F, et al. Voluntary exercise increases oligodendrogenesis in spinal cord. Int J Neurosci. 2010;120(4):280–90.

22. Dougherty KD, Dreyfus CF, Black IB. Brain-derived neurotrophic factor in astrocytes, oligodendrocytes, and microglia/macrophages after spinal cord injury. Neurobiol Dis. 2000;7(6 Pt B):574–85.

23. Asano K, Nakamura T, Funakoshi K. Early mobilization in spinal cord injury promotes changes in microglial dynamics and recovery of motor function. IBRO Neurosci Rep. 2022;12:366–76.

24. Bai J, Geng B, Wang X, Wang S, Yi Q, Tang Y, et al. Exercise Facilitates the M1-to-M2 Polarization of Microglia by Enhancing Autophagy via the BDNF/AKT/mTOR Pathway in Neuropathic Pain. Pain Physician. 2022;25(7):E1137–E51.

25. Chhaya SJ, Quiros-Molina D, Tamashiro-Orrego AD, Houle JD, Detloff MR. Exercise-Induced Changes to the Macrophage Response in the Dorsal Root Ganglia Prevent Neuropathic Pain after Spinal Cord Injury. J Neurotrauma. 2019;36(6):877–90.

26. Wang MJ, Jing XY, Wang YZ, Yang BR, Lu Q, Hu H, et al. Exercise, Spinal Microglia and Neuropathic Pain: Potential Molecular Mechanisms. Neurochem Res. 2024;49(1):29–37.

27. Brennan FH, Li Y, Wang C, Ma A, Guo Q, Li Y, et al. Microglia coordinate cellular interactions during spinal cord repair in mice. Nat Commun. 2022;13(1):4096.

28. Belaya I, Ivanova M, Sorvari A, Ilicic M, Loppi S, Koivisto H, et al. Astrocyte remodeling in the beneficial effects of long-term voluntary exercise in Alzheimer’s disease. J Neuroinflammation. 2020;17(1):271.

29. Tsai SF, Chen PC, Calkins MJ, Wu SY, Kuo YM. Exercise Counteracts Aging-Related Memory Impairment: A Potential Role for the Astrocytic Metabolic Shuttle. Front Aging Neurosci. 2016;8:57.

30. Matsui T, Omuro H, Liu YF, Soya M, Shima T, McEwen BS, et al. Astrocytic glycogen-derived lactate fuels the brain during exhaustive exercise to maintain endurance capacity. Proc Natl Acad Sci U S A. 2017;114(24):6358–63.

31. Li F, Geng X, Yun HJ, Haddad Y, Chen Y, Ding Y. Neuroplastic Effect of Exercise Through Astrocytes Activation and Cellular Crosstalk. Aging Dis. 2021;12(7):1644–57.

32. Maugeri G, D’Agata V, Magri B, Roggio F, Castorina A, Ravalli S, et al. Neuroprotective Effects of Physical Activity via the Adaptation of Astrocytes. Cells. 2021;10(6).

33. Saur L, Baptista PP, de Senna PN, Paim MF, do Nascimento P, Ilha J, et al. Physical exercise increases GFAP expression and induces morphological changes in hippocampal astrocytes. Brain Struct Funct. 2014;219(1):293–302.

34. Ying X, Yu X, Zhu J, Li X, Zheng Y, Xie Q, et al. Water Treadmill Training Ameliorates Neurite Outgrowth Inhibition Associated with NGR/RhoA/ROCK by Inhibiting Astrocyte Activation following Spinal Cord Injury. Oxid Med Cell Longev. 2022;2022:1724362.

35. Chen Y, Wei Y, Liu J, Zhu T, Zhou C, Zhang D. Spatial transcriptomics combined with single-nucleus RNA sequencing reveals glial cell heterogeneity in the human spinal cord. Neural Regen Res. 2025;20(11):3302–16.

36. Matson KJE, Russ DE, Kathe C, Hua I, Maric D, Ding Y, et al. Single cell atlas of spinal cord injury in mice reveals a pro-regenerative signature in spinocerebellar neurons. Nat Commun. 2022;13(1):5628.

37. Russ DE, Cross RBP, Li L, Koch SC, Matson KJE, Yadav A, et al. A harmonized atlas of mouse spinal cord cell types and their spatial organization. Nat Commun. 2021;12(1):5722.

38. Milich LM, Choi JS, Ryan C, Cerqueira SR, Benavides S, Yahn SL, et al. Single-cell analysis of the cellular heterogeneity and interactions in the injured mouse spinal cord. J Exp Med. 2021;218(8).

39. Skinnider MA, Gautier M, Teo AYY, Kathe C, Hutson TH, Laskaratos A, et al. Single-cell and spatial atlases of spinal cord injury in the Tabulae Paralytica. Nature. 2024;631(8019):150–63.

40. Li C, Wu Z, Zhou L, Shao J, Hu X, Xu W, et al. Temporal and spatial cellular and molecular pathological alterations with single-cell resolution in the adult spinal cord after injury. Signal Transduct Target Ther. 2022;7(1):65.

41. Blum JA, Klemm S, Shadrach JL, Guttenplan KA, Nakayama L, Kathiria A, et al. Single-cell transcriptomic analysis of the adult mouse spinal cord reveals molecular diversity of autonomic and skeletal motor neurons. Nat Neurosci. 2021;24(4):572–83.

42. Floriddia EM, Lourenco T, Zhang S, van Bruggen D, Hilscher MM, Kukanja P, et al. Distinct oligodendrocyte populations have spatial preference and different responses to spinal cord injury. Nat Commun. 2020;11(1):5860.

43. Zhang D, Chen Y, Wei Y, Chen H, Wu Y, Wu L, et al. Spatial transcriptomics and single-nucleus RNA sequencing reveal a transcriptomic atlas of adult human spinal cord. Elife. 2024;12.

44. Yadav A, Matson KJE, Li L, Hua I, Petrescu J, Kang K, et al. A cellular taxonomy of the adult human spinal cord. Neuron. 2023;111(3):328–44 e7.

45. Sun S, Ma S, Cai Y, Wang S, Ren J, Yang Y, et al. A single-cell transcriptomic atlas of exercise-induced anti-inflammatory and geroprotective effects across the body. Innovation (Camb). 2023;4(1):100380.

46. Du J, Hou J, Du N, Zhang X. Single-Nucleus RNA-Seq Reveals Aerobic Exercise-Induced Remodeling of Spinal Cord Aging in Mice. FASEB J. 2025;39(23):e71205.

47. Xin W, Chan JR. Myelin plasticity: sculpting circuits in learning and memory. Nat Rev Neurosci. 2020;21(12):682–94.

48. Fang LP, Bai X. Oligodendrocyte precursor cells: the multitaskers in the brain. Pflugers Arch. 2023;475(9):1035–44.

49. Jin S, Plikus MV, Nie Q. CellChat for systematic analysis of cell-cell communication from single-cell transcriptomics. Nat Protoc. 2025;20(1):180–219.

50. Houle JD, Cote MP. Axon regeneration and exercise-dependent plasticity after spinal cord injury. Ann N Y Acad Sci. 2013;1279(1):154–63.

51. Lynskey JV, Belanger A, Jung R. Activity-dependent plasticity in spinal cord injury. J Rehabil Res Dev. 2008;45(2):229–40.

52. Sancho L, Contreras M, Allen NJ. Glia as sculptors of synaptic plasticity. Neurosci Res. 2021;167:17–29.

53. Lago-Baldaia I, Fernandes VM, Ackerman SD. More Than Mortar: Glia as Architects of Nervous System Development and Disease. Front Cell Dev Biol. 2020;8:611269.

54. Allen NJ, Lyons DA. Glia as architects of central nervous system formation and function. Science. 2018;362(6411):181–5.

55. Saab AS, Tzvetanova ID, Nave KA. The role of myelin and oligodendrocytes in axonal energy metabolism. Curr Opin Neurobiol. 2013;23(6):1065–72.

56. Ioannou MS, Jackson J, Sheu SH, Chang CL, Weigel AV, Liu H, et al. Neuron-Astrocyte Metabolic Coupling Protects against Activity-Induced Fatty Acid Toxicity. Cell. 2019;177(6):1522–35 e14.

57. Gibson EM, Purger D, Mount CW, Goldstein AK, Lin GL, Wood LS, et al. Neuronal activity promotes oligodendrogenesis and adaptive myelination in the mammalian brain. Science. 2014;344(6183):1252304.

58. Mensch S, Baraban M, Almeida R, Czopka T, Ausborn J, El Manira A, et al. Synaptic vesicle release regulates myelin sheath number of individual oligodendrocytes in vivo. Nat Neurosci. 2015;18(5):628–30.

59. Vukovic J, Colditz MJ, Blackmore DG, Ruitenberg MJ, Bartlett PF. Microglia modulate hippocampal neural precursor activity in response to exercise and aging. J Neurosci. 2012;32(19):6435–43.

60. Mee-Inta O, Zhao ZW, Kuo YM. Physical Exercise Inhibits Inflammation and Microglial Activation. Cells. 2019;8(7).

61. Strohm AO, Majewska AK. Physical exercise regulates microglia in health and disease. Front Neurosci. 2024;18:1420322.

62. van Praag H, Christie BR, Sejnowski TJ, Gage FH. Running enhances neurogenesis, learning, and long-term potentiation in mice. Proc Natl Acad Sci U S A. 1999;96(23):13427–31.

63. Morland C, Andersson KA, Haugen OP, Hadzic A, Kleppa L, Gille A, et al. Exercise induces cerebral VEGF and angiogenesis via the lactate receptor HCAR1. Nat Commun. 2017;8:15557.

64. Swain RA, Harris AB, Wiener EC, Dutka MV, Morris HD, Theien BE, et al. Prolonged exercise induces angiogenesis and increases cerebral blood volume in primary motor cortex of the rat. Neuroscience. 2003;117(4):1037–46.

65. Alkaslasi MR, Piccus ZE, Hareendran S, Silberberg H, Chen L, Zhang Y, et al. Single nucleus RNA-sequencing defines unexpected diversity of cholinergic neuron types in the adult mouse spinal cord. Nat Commun. 2021;12(1):2471.

66. Furrer R, Hawley JA, Handschin C. The molecular athlete: exercise physiology from mechanisms to medals. Physiol Rev. 2023;103(3):1693–787.

67. Matson KJE, Sathyamurthy A, Johnson KR, Kelly MC, Kelley MW, Levine AJ. Isolation of Adult Spinal Cord Nuclei for Massively Parallel Single-nucleus RNA Sequencing. J Vis Exp. 2018(140).

68. Fleming SJ, Chaffin MD, Arduini A, Akkad AD, Banks E, Marioni JC, et al. Unsupervised removal of systematic background noise from droplet-based single-cell experiments using CellBender. Nat Methods. 2023;20(9):1323–35.

69. Hao Y, Stuart T, Kowalski MH, Choudhary S, Hoffman P, Hartman A, et al. Dictionary learning for integrative, multimodal and scalable single-cell analysis. Nat Biotechnol. 2024;42(2):293–304.

70. Love MI, Huber W, Anders S. Moderated estimation of fold change and dispersion for RNA-seq data with DESeq2. Genome Biol. 2014;15(12):550.

71. Zhou Y, Zhou B, Pache L, Chang M, Khodabakhshi AH, Tanaseichuk O, et al. Metascape provides a biologist-oriented resource for the analysis of systems-level datasets. Nat Commun. 2019;10(1):1523.

